# De novo protein ligand design including protein flexibility and conformational adaptation

**DOI:** 10.64898/2026.01.08.698398

**Authors:** Jakob Agamia, Martin Zacharias

## Abstract

**Motivation:** The rational design of chemical compounds that bind to a desired protein target molecule is a major goal of drug discovery. Most current molecular docking but also fragment-based build-up or machine-learning based generative drug design approaches employ a rigid protein target structure.

**Results:** Based on recent progress in predicting protein structures and complexes with chemical compounds we have designed an approach, AI-MCLig, to optimize a chemical compound bound to a fully flexible and conformationally adaptable protein binding region. During a Monte-Carlo (MC) type simulation to randomly change a chemical compound the target protein-compound complex is completely rebuilt at every MC step using the Chai-1 protein structure prediction program. Besides compound flexibility it allows the protein to adapt to the chemically changing compound. MC-protocols based on atom/bond type changes or based on combining larger chemical fragments have been tested. Simulations on three test targets resulted in potential ligands that show very good binding scores comparable to experimentally known binders using several different scoring schemes. The MC-based compound design approach is complementary to existing approaches and could help for the rapid design of putative binders including induced fit of the protein target.

**Availability and implementation:** Datasets, examples and source code are available on our public GitHub repository https://github.com/JakobAgamia/AI-MCLig and on Zenodo at https://doi.org/10.5281/zenodo.17800140.

## 1 Introduction

Protein molecules and complexes formed between proteins and other types of biomolecules play key roles in basically all biological processes [1]. Many drug design efforts aim at identifying or developing small organic molecules that specifically bind to target proteins and interfere with its function [2, 3, 4]. Such ligands can potentially inhibit protein function or influence interactions with biomolecule partners [5].

Computational approaches play an increasing role to screen for ligands that can bind to protein pockets with high affinity. Traditionally, in silico pharmacophore screening and molecular docking methods are used to select putative binders from large data bases [6, 7, 8, 3]. Screening of large data bases with multi-billion of compounds [9, 10] is computationally demanding and faces the difficulty to accurately score putative binders [11, 10, 12]. An additional difficulty arises from the fact that most virtual screening methods employ rigid protein target structures, completely neglecting even small adaptations of the protein structure upon ligand binding[13, 3]. In recent years artificial intelligence (AI) driven methods have gained popularity and allow the generation of entirely new compounds in de novo drug design[6, 12, 10, 14, 7]. Aim of de novo compound design methods is to identify or newly generate chemical compounds (generative models) that fit to a given target protein pocket and bind with high-affinity. A variety of AI-based methods are available ranging from generative adversial networks (GANs), variational auto-encoders (VAEs) and more recently diffusion models (e.g. reviewed in [14]). The methods are trained on large databases of known protein-ligand complexes[15, 16].

Typically, such approaches employ a defined and fixed (rigid) target pocket and can generate entirely new potential binders thereby including also biases for solubility and chemical accessibility of the generated compounds[4, 17]. However, one major drawback of both traditional molecular docking and generative models is the typical assumption of a rigid protein target pocket that realistically may change shape and adapt to some degree to a bound ligand[13, 3].

AI-based structure prediction approaches such as AlphaFold2 (AF2)[18] allow the accurate three-dimensional (3D) modeling of proteins and protein complexes. More recently, the deep learning methodology was extended to not only accurately model protein structures but also complexes with organic molecules using AlphaFold3 (AF3)[19] and related programs such as Boltz1[20] and Chai-1[21]. In particular, Chai-1 allows quite rapid generation of protein three dimensional (3D) structures in complex with organic drug-like ligands.

Based on Chai-1 we have developed Monte-Carlo (MC) type simulation approaches in chemical space (termed AI-MCLig) to generate new organic compounds that potentially bind at a protein target region. The first protocol uses a set of atomistic (e.g. atom-type, bond-type) chemical changes to modify a simple starting structure (e.g. benzene). The second method recombines molecular fragments based on the BRICS [22] method. We demonstrate that the AI-MCLig approach can recover a desired ligand if structural similarity is used as target score. In case of de novo ligand generation for a desired protein target, for both AI-MCLig protocols the ligand-protein complex is completely rebuilt (using the artificial intelligence, AI-based, Chai-1 method at each MC step). Besides of including compound flexibility it offers an important additional advantage of the method allowing for full conformational adaptation of the protein structure upon ligand modification. Whether or not the change is accepted, is based mostly on the change in the Chai-1 confidence score. On three examples we demonstrate that the MC method allows to rapidly generate new putative binders with favorable scoring including also alternative force field based scoring. The method could complement traditional molecular docking as well as generative de novo drug design approaches that mostly employ rigid binding pockets.

## 2 Methods

### 2.1 Monte Carlo simulation approach

In the Monte Carlo (MC) simulation, the ligand is built by modifying a current compound structure with elementary chemical steps (see below). The modifications are introduced using the rdkit package[23] which includes routines to ensure the chemical validity of the modified compound. The compound chemical smiles [24] and the protein sequence are used with Chai-1[21] for AI-based rebuilding and evaluation (termed AI-MCLig approach). The scoring includes mostly the confidence score, a synthetic accessibility bias, a solubility bias and a drug-likeness bias (see paragraph below). The newly generated ligand structure and placement is accepted if the score improves, otherwise the structure is only accepted with a probability of

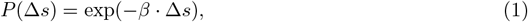

where Δ*s* is the difference between the new score and current score and *β* is a factor to influence the probability of accepting a less favorable structure in the simulation. It corresponds to the inverse of an effective temperature of the MC type simulation (small *β* translates to a broad sampling of compounds with various scores and high *β* is a very selective search accepting mostly well scoring compounds).

The chemical steps used by the simulation are:

**Adding small chemical groups (standard probability 0.3)** In this step a small chemical group (e.g. CH3, OH) is added at a randomly chosen accessible point in the molecule replacing another group (e.g. H-atom).

**Removing atoms (standard probability 0.2)** A randomly chosen atom in the molecule is removed.

**Changing atom types (standard probability 0.1)** The type of a randomly chosen atom is changed to a different type.

**Adding atoms in chain (standard probability 0.15)** An atom is added in between two atoms. While this step is not strictly necessary to reach all chemical structures, it helps to avoid local minima.

**Changing bond types (standard probability 0.025)** A randomly chosen bond is changed from single to double or vice versa.

**Forming rings (standard probability 0.025)** Chains of sufficient length to form a ring are identified. One of those is randomly chosen and a bond is formed between two atoms to form a ring structure.

**Breaking up rings (standard probability 0.05)** A random bond in a ring is selected and removed to break up a ring. If the ring was aromatic, the involved atoms and bonds are automatically set to non-aromatic.

**Turning rings aromatic (standard probability 0.05)** A randomly chosen non-aromatic ring is turned aromatic or vice-versa.

**Rearranging bonds (standard probability 0.1)** In this step an atom with more than two bonds is selected. One of the bonds is removed and reformed with a neighboring atom. This step is again not strictly necessary but helpful in avoiding local minima.

Which step is used each time is randomly chosen with probabilities for chemical changes as given above. The probabilities for each step were determined by testing different probability sets on three targets (bromodomain, p38 map kinase and Pim1 kinase). The set of probabilities used in the MC search correspond to the most efficient combination (see figure SI 1 for comparison with other probabilities). It should be mentioned, that since the simulations are time consuming the number of tested parameter sets is limited. Hence, optimal parameter sets might also vary for different protein targets which will be investigated in future studies. Each one-hundredth step the structure is reset to the best structure found up to that point avoiding local minima. We term this type of AI-MCLig simulation atomistic-step MC simulation.

### 2.2 Fragment-based MC simulation

The fragment-based MC simulation follows the same concept, however, instead of using basic chemical steps, the ligand is built by recombining molecular fragments based on rdkit’s[23] implementation of the BRICS[22] method. During the simulation entire chemical fragments are added or removed from the molecule using the same criteria as in the atomistic-step MC procedure. To determine which fragments are compatible with each other, the rules from rdkit’s[23] implementation of BRICS were followed. The two steps used by this simulation are:

**Adding fragments** A placeholder in the molecule is chosen at random and an appropriate fragment is added at that position.

**Removing fragments** A random bond between fragments is chosen. It is removed and replaced with the placeholder, which occupied that position before the bond.

### 2.3 Scoring of compounds during MC simulations

As mentioned before, the score for whether a change is kept or not is mostly based on the Chai-1 confidence score. However three biases are introduced to the score to enforce desired properties:

**Synthetic accessibility(SA) bias** The ligands resulting from the simulations should not only have good binding affinities, but also be synthesized with reasonable effort. Therefore a bias with the SA score[25] is added. It should be noted that for theses simulations the SA score is considered maximal below a threshold. This is because attempting to enforce a very low SA score is unreasonable, as the ligand must have a certain amount of complexity to fit the binding site.

**Solubility bias** To ensure the resulting molecules are soluble in water, a bias with the ESOL score[26] is added.

**Drug-likeness bias** The ligands should ideally have drug-like properties. Therefore rdkit’s quantitative estimate of drug-likeness (QED) score, which is based on [27], is added (neglecting molecular weight factor).

With these biases the final score used for all simulations is derived as

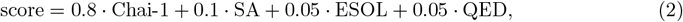

with the SA and ESOL score being normalized to also lie between zero and one. The contribution of the Chai-1 score is set to 0.8, as it has been found that a simulation with a smaller contribution often fails to maximize this score. The contribution of the SA score should only be high enough to hold the simulation under the given threshold. A value of 0.1 has been found to do this in almost all cases. The QED and ESOL scores only make small contributions. However, even with these low scoring contributions, we found that the QED and ESOL scores of the generated compounds remain in a reasonable range for most of the tested cases and still allow generation of a large variety of compounds. In the Supplementary Material (see Results section) simulations with a variation of score compositions are included demonstrating the deviations from our combination indicated above generally lead to worse results.

### 2.4 Molecular Dynamics simulations and MMGBSA calculations

Some of the resulting ligands were additionally evaluated with Molecular Dynamics (MD) simulations and MMGBSA (Molecular Mechanics Generalized Born Surface Area) calculations[28]. The initial coordinate files were generated using Chai-1[21]. Ligand force field parameters were obtained using the antechamber program [29, 30]. The tleap module of the Amber package [31, 32] was used to solvate the protein-ligand complex using TIP3P water model [33]. The complex was energy minimized (2000 steps) and then heated within 1 ns to a temperature of 300 K including positional restraints on each non-hydrogen atom. Subsequently, an 20 ns unrestraint MD simulation, using pmemd.cuda [31, 32, 34, 35, 36] was performed. The first 2.5*ns* of the trajectory were disregarded. The remaining trajectory (750 frames) was then processed by the MMGBSA tool of the Amber package (using igb=5) [32, 28]. The MDAnalysis package [37, 38, 39, 40, 41] was used for calculation of root-mean-square deviations (RMSD).

## 3 Results and discussion

### 3.1 MC based recovery of target ligands from simple starting compounds

For our MC type searches in chemical space it is important to assess, whether relevant chemical structures can in principal be reached using the employed MC steps. To this end MC simulations were performed with the regular score being replaced with a dice similarity score, derived with rdkit[23]. The dice score is calculated from atom-pair fingerprints [42], as it has been found that this type of fingerprints can be calculated very efficiently in our MC simulations. The dice score was selected for this, because it functioned somewhat better with the simulation than similar alternatives, which were found to be even more prone to local minima. The similarity was calculated between the current compound and a known target compound. The target compounds were taken from [16], mainly from complexes of the bromodomain, serine/threonine-protein kinase pim-1 and p38 map kinase. The same targets were subsequently also used for the design of new putative binders. Note, the dice similarity score for a compound with respect to a reference compound represents a very rough scoring landscape in chemical space with many local minima and barriers. Therefore, the atom-based simulations consisted of two stages. For each case a simple benzene molecule was used as a starting compound. In the first stage 30 simulations were run simultaneously for 5000 steps. Every 500 steps the structure of all 30 simulations was set to the best structure found by all simulations. The second stage consisted of a single simulation with 10000 MC steps and reset to the best structure found at every 100th step. The *β* factor was set to 50 for all simulations to provide high selectivity. Its leads to a reasonable acceptance rate of 0.1-0.2 to select efficiently among a very large number of possible chemical changes those few that lead in the right direction of similarity to the target compound.

A score of 1 indicates identity between MC generated structure and target structure (see SI Fig. SI 2 and Fig. SI 3). Especially for ligands with many ring structures it is often very difficult to escape out of certain local minima. Nevertheless, the results indicate (Table 1) that for most cases the approach succeeds in producing exactly the desired target compound. For some of the more complicated compounds the simulation got stuck in local minima that differ from the target reference compound. Nevertheless, it indicates that even on a very rough score landscape the MC procedure can reach or comes very close to a desired target chemical structure. The probabilities for different types of chemical changes for each of the MC steps are shown in section 2.1. Variations of these step parameters and of the effective simulation temperature were tested (SI Fig. SI 4, Table SI 1, Table SI 2) but resulted in reduced performance compared to the standard parameters (Methods).

**Table 1:**
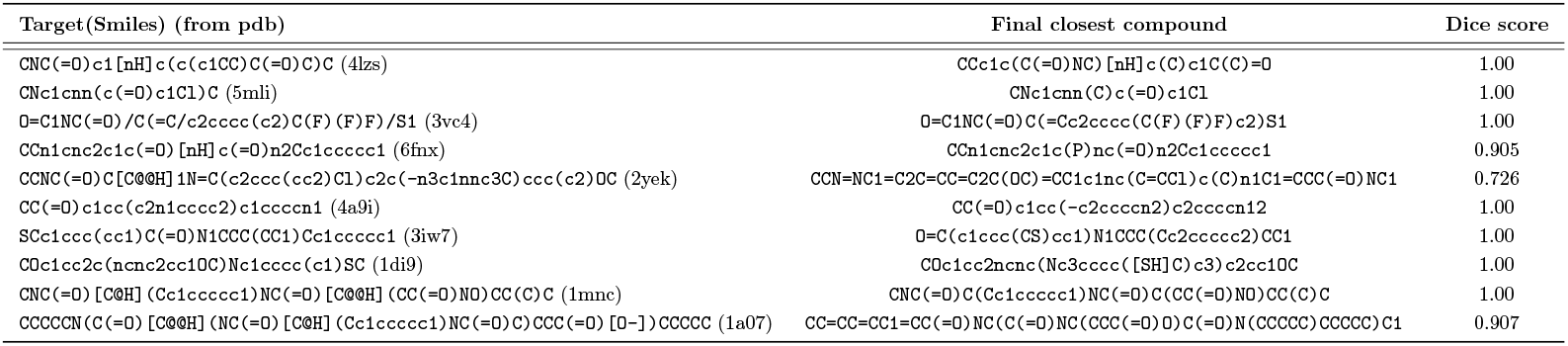
The target and resulting compounds in SMILES representation for atomistic-step MC simulations. Benzene was used as starting compound in the atomistic-step MC approach and the dice similarity score was optimized in the MC search. The target SMILES correspond to bromodomain ligands extract from PDB-bind [16].

For the MC search approach based on combining chemical fragments, the known ligand compound was broken down into fragments used in a simulation of 10000 steps. Since the addition or removal of whole chemical fragments causes larger dice score changes compared to the atomistic-step MC approach a smaller *β* factor of 5 was necessary to achieve a reasonable MC acceptance rate. As fragments can only be removed or added, often reaching a new structure may require acceptance of intermediates with a significantly reduced score. The standard probabilities for each type of step were 0.6 for adding fragments and 0.4 for removing fragments. The structure was reset to the best structure found at every 50th step. A random fragment of the target ligand was used as starting structure. All tested ligand compounds were successfully reassembled (SI Table SI 3). Examples for the dice score vs. MC steps are given in the SI (SI Fig. SI 5 including different choices of *β* and probability parameters for adding or removing fragments).

### 3.2 Correlation of Chai-1 confidence score and ligand binding affinity

Similar to AF3 or Boltz1, the Chai-1 structure prediction program provides a pLDDT confidence score that is typically a very good measure for the reliability of a predicted protein structure. Such confidence score is also provided in case of a protein-ligand complex. However, it is not clear how well it correlates with experimental binding affinities for known complexes. We selected three protein cases for which structures of many complexes and corresponding binding affinities are available. To this end, the protein-ligand complexes for the bromodomain, serine/threonine-protein kinase pim-1 and p38 map kinase were evaluated for a number of different ligands (data taken from PDBbind+ data set[16]). The Chai-1 scores were compared to the known binding affinities (Fig. 1). While there is no direct correlation, a very high Chai-1 score does typically correspond to a high average binding affinity. Although the Chai-1 confidence score is far from ideal the results suggest, however, that maximizing the Chai-1 score is a reasonable approach for finding ligands with potentially high binding affinities. It should be noted, that the correlation depends on the protein target and might be worse for the different cases, than the ones shown here.

**Figure 1.**
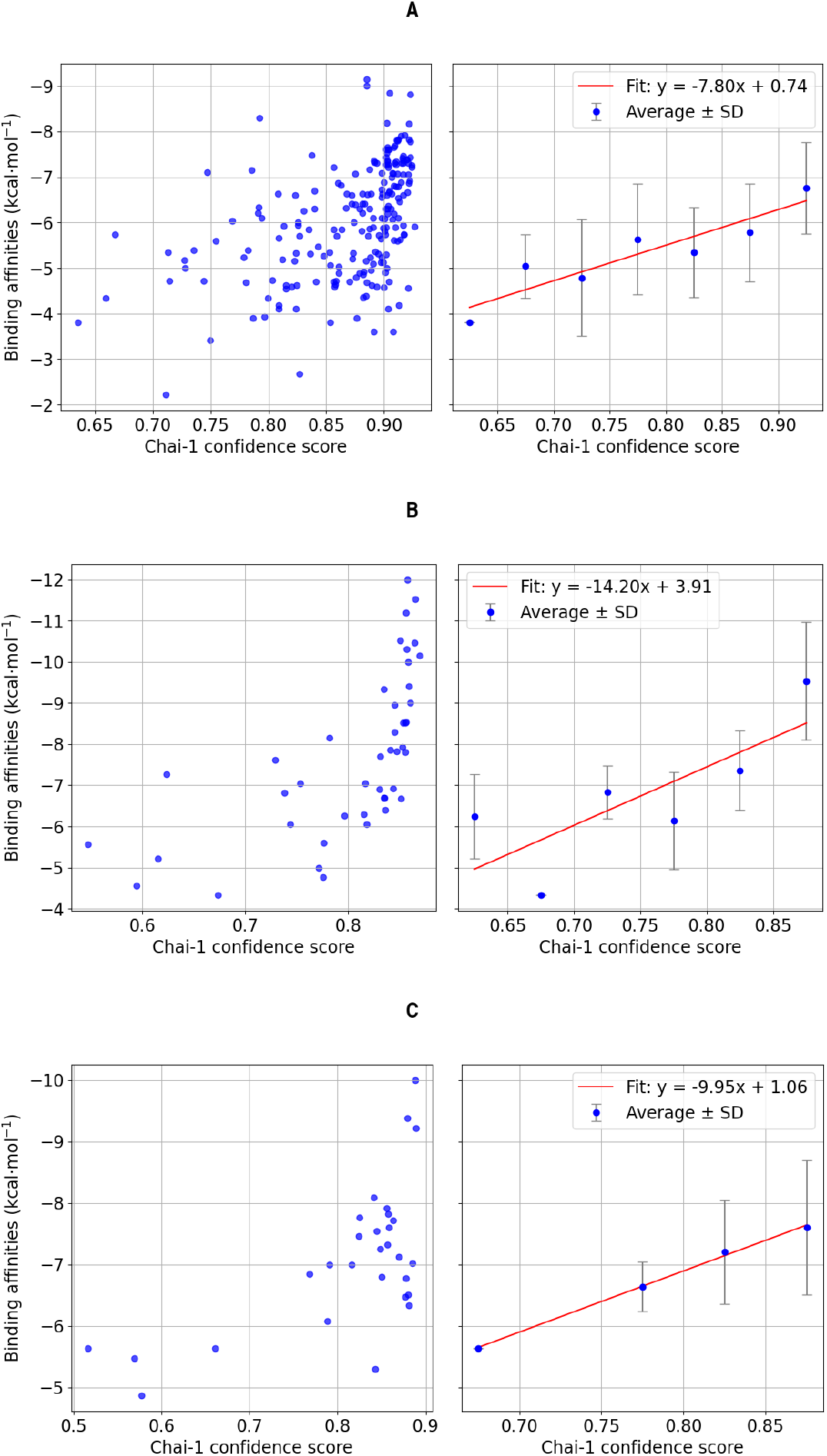
Illustration of the correlation between the experimental ligand binding affinity and the Chai-1 confidence score. (A) Experimental binding affinity against the Chai-1 confidence score for different ligands of the bromodomain complex. (B) Experimental binding affinity against the Chai-1 confidence score for ligands of the serine/threonine-protein kinase pim-1 complex. (C) Binding affinity against the Chai-1 confidence score for ligands of the p38 map kinase complex.

Nevertheless, it is also important to realize that the realistic scoring of protein-ligand complexes is still a major challenge and needs further improvement in future work. We also investigated the possibility to use other more force field based scoring provided by the Autodock Vina docking approach [43], [44] as an alternative to the Chai-1 confidence score. The correlation between the Autodock score and the binding affinity was similarly tested (SI Fig. SI 6) and does not appear to be much better than that of the Chai-1 score and when used in simulations the Chai-1 score generally produced more promising results than the auto dock score. Efforts to combine the Autodock and Chai-1 scores (as a sum of normalized Autodock score and Chai-1 score) did not result in improved correlation (SI Fig. SI 7). Hence, although not ideal the Chai-1 confidence score (in combination with additional contributions, see Methods) was used for all following simulations.

### 3.3 De novo generation of bound ligands using atomistic-step MC simulation

The AI-MCLig approach based on atomic changes of bound ligands was applied to the bromodomain, the serine/threonine-protein kinase pim-1 protein, the p38 map kinase protein and the *β*-1 adrenergic receptor (a seven helix bundle GPCR protein). Each simulation consisted of 2000 MC steps, with a constant *β* = 50 and starting from benzene molecules. The probabilities for each type of changes in a MC step are the same as for the similarity based simulation. For each protein complex we performed 3 MC simulations using different initial random seeds. Each MC run took roughly 20 hours, slightly depending on protein size, on a PC with NVIDIA RTX4090 card.

In order to illustrate the progressive chemical changes of the ligand structure, every 20th accepted structure of the first 100 MC steps of the first simulation on the bromodomain complex are shown in Fig. 2. It also indicates how the MC approach progressively fills the protein pocket with sterically well fitting ligands.

**Figure 2.**
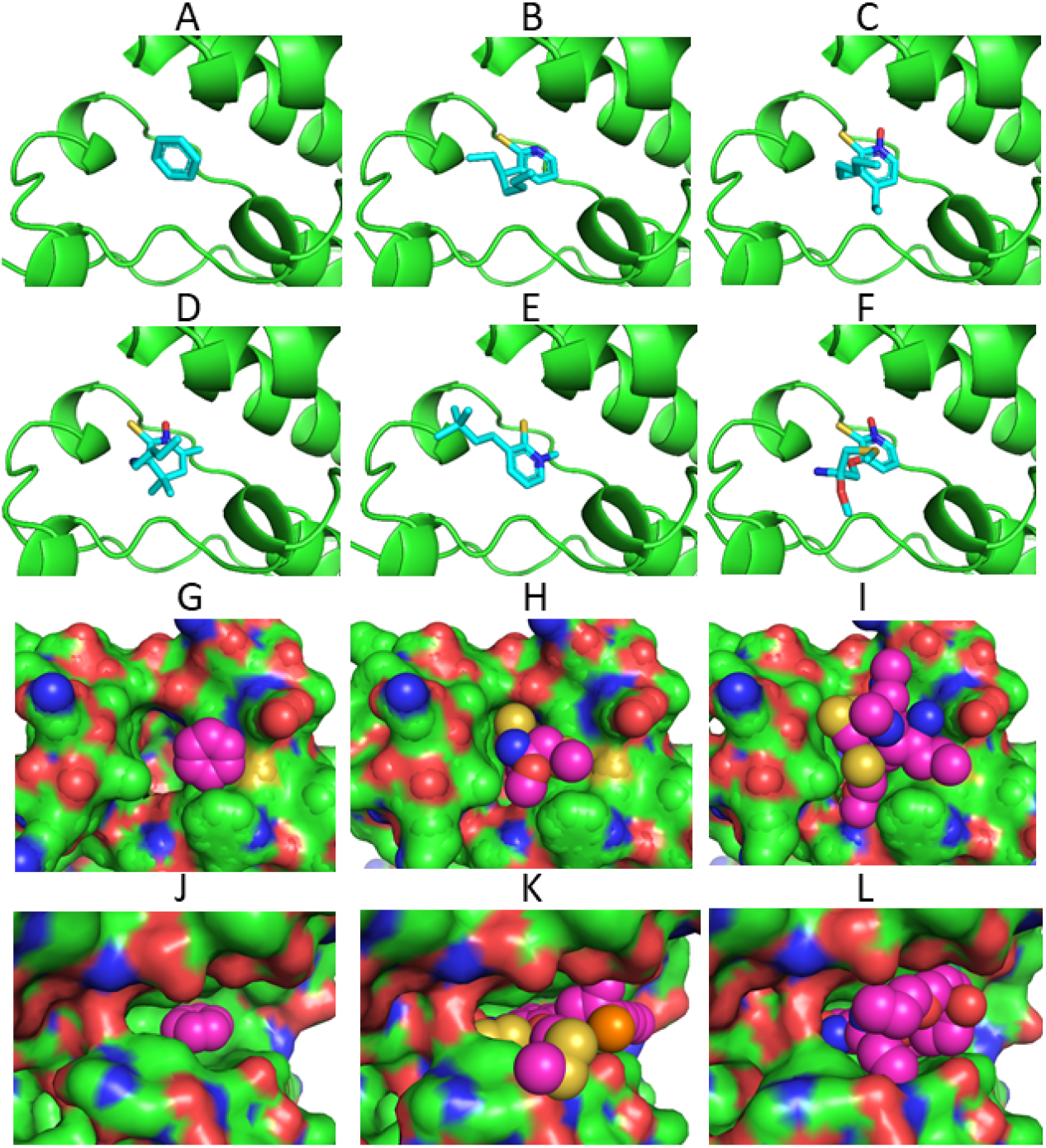
Illustration of the MC-based ligand design process. (A-F) Generated ligands (blue stick model) after 0, 20, 40, 60, 80 and 100 MC steps for the bromodomain ligand binding pocket (protein as green cartoon). (G) Benzene (pink spheres) start ligand placed in the bromodomain binding pocket. (H) same as (G) with the generated ligand after 200 MC steps. (I) same as (G) but after 2000 MC steps. Note, the generated ligand tightly fits into the binding pocket (protein represented as solvent accessible surface). (J-L) same as (G-I) but for the serine/threonine pim-1 kinase target.

In each case the finally generated compounds are bound at the target protein in the desired binding pocket and result in a high Chai-1 confidence score (Table 2). Since the MC type simulation is a stochastic search method it is not unexpected that the final compounds differ for each MC run. It is also compatible with the experimental observation that indeed many chemically distinct ligands can be found for the four target proteins that bind with nano- or micromolar affinity to the proteins. Encouragingly, the high final Chai-1 scores are also comparable to the scores for known ligands for each target that bind experimentally with highest affinity (compare Table 2 and Table 3). As explained in the previous section, this should correspond, at least on average, to a very favorable binding affinity. During the MC run the score improves initially within the first 250 steps quite rapidly and evolves further more gradually (illustrated in Fig. 3, see also SI Fig. SI 8). After the initial phase, improvements that are simultaneously chemically feasible, improve interactions and allow good packing without sterical overlap become increasingly difficult for the MC procedure. In the future, it might be possible to design smarter MC moves in chemical space to increase the probability of improvement at each MC step.

**Table 2:**
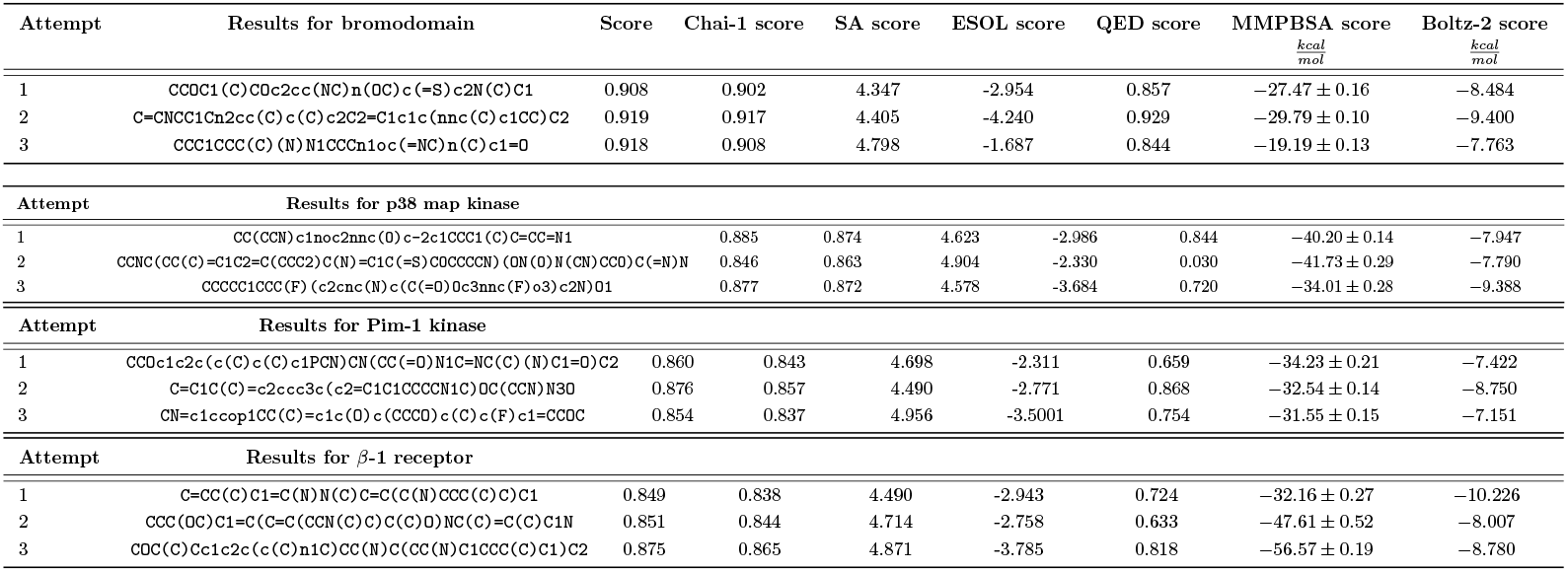
Final generated compounds (Smiles code, some of the Smiles are split into two lines) resulting from the atomistic-step MC method for the bromodomain, p38 kinase and serine/threonine-protein kinase pim-1 and the final scores.

**Table 3:**
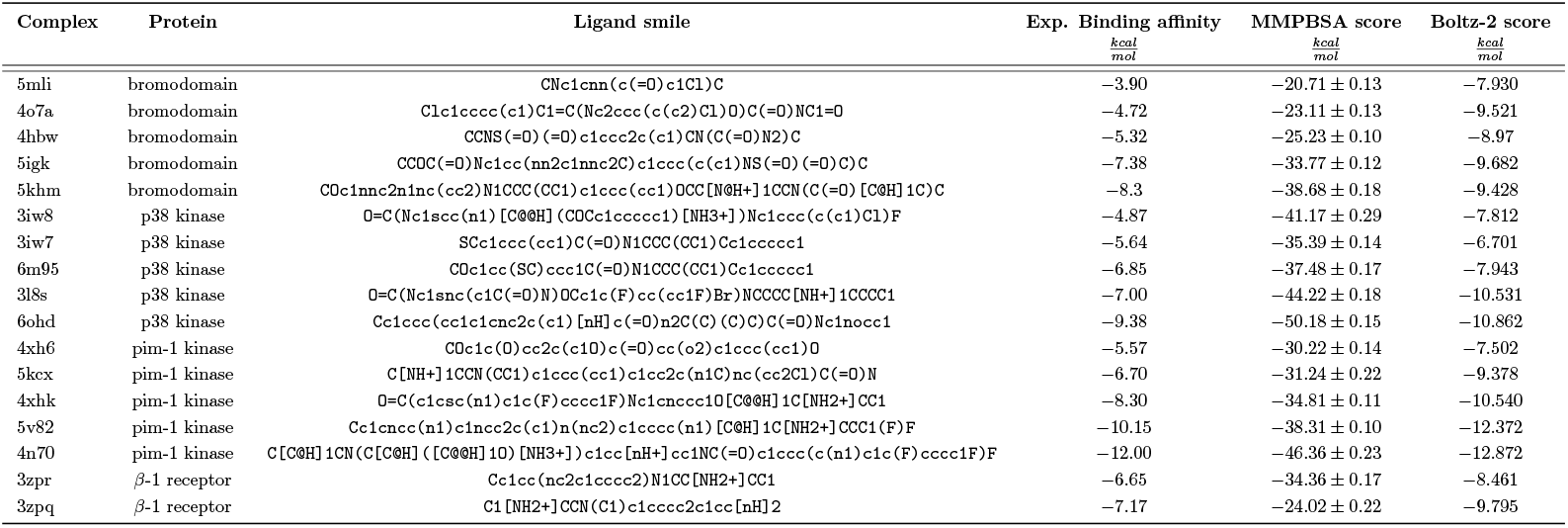
Experimental binding affinities, MMGBSA score and Boltz-2 score for ligands of the bromod-omain, the serine/threonine-protein kinase pim-1 protein and the p38 map kinase protein.

**Figure 3.**
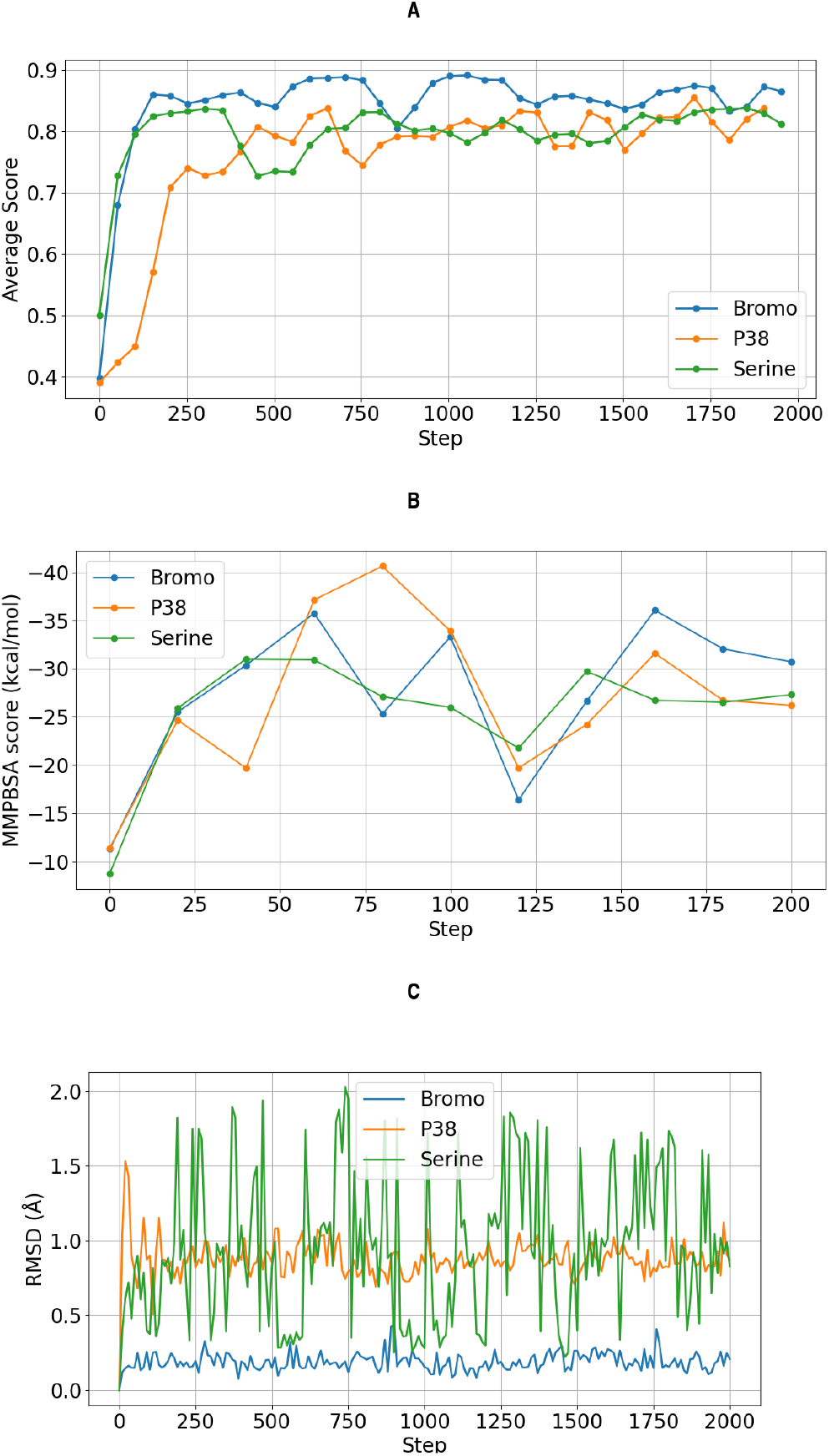
(A) The average of the score for every fifty steps during the basic MC simulation for one design run on each of the three protein targets. The average was taken as running windows over 100 MC steps. (B) Calculated MMGBSA score of 3 MC simulation runs for the first 200 MC steps. (C) Root-mean-square deviation (RMSD) of the non-hydrogen protein atoms forming the ligand binding pocket with respect to the start structure. The pocket protein atoms are defined as all atoms within 5 of the native ligand

In the previous paragraph we found only an average correlation of the Chai-1 score and the experimental ligand binding affinity for the test proteins. Hence, it is important to evaluate the generated compounds using alternative force field based approaches. We used the widely employed MMGBSA (Molecular Mechanics Generalize Born) method [28] to estimate the mean interaction between compounds and target proteins. In an evaluation of a large number of protein-ligand complexes [11] mean MMGBSA interaction energies of 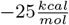 to 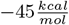 were typically found for high-affinity binders. The MMGBSA results for the 3 runs and for each protein target case are in this range (Table 2) and encouragingly are also of similar magnitude as MMGBSA results for experimentally evaluated protein binders (Table 3). Note also, that the MMGBSA scores for the four test proteins and different ligands correlate qualitatively with experimentally known binding affinity (Table 3).

We also re-calculated the MMGBSA score for the first 200 MC steps of one simulation for each target (Fig. SI 9) and the calculated averages of the 3 runs on each protein (Fig. 3). Similar to the Chai-1 score it improves initially quite rapidly but subsequently only gradually.

As another independent test the generated compounds were also evaluated with the recent ligand-receptor binding affinity model of Boltz-2 [45]. This model returns a log(IC50) score for the evaluation of ligand-receptor complexes and the accuracy of the model is considered to be similar to alchemical absolute binding free energies that are considered as most accurate binding prediction method [45]. The log(IC50) score was converted into a pIC50 in kcal/mol. Also, the Boltz-2 score assigns quite favorable predicted binding scores to the MC generated compounds (Table 2). One important advantage of the present MC based ligand generation method is the full reconstruction of the protein partner allowing for conformational induced fit adaptation of the protein. The variation of the protein structure during MC based ligand generation depends significantly on the protein target (Fig. 3, see also Fig. SI 10. No major structural changes were observed for the bromodomain protein. It indicates a relatively rigid binding pocket which varies little in structure upon binding different ligands. Indeed, this finding compares well with experimental structures of the bromodomain that change only little in complexes with different ligands (Fig. SI 11). It is important to note that even small changes in conformation can be of critical importance for correctly placing and evaluation of protein-ligand complexes (a major problem of rigid protein-ligand docking approaches).

In contrast, the structural variations during the MC simulations are larger for the serin kinase and P38 cases (Fig. 3). This is also in line with the experimentally found larger structural variation in complex with different ligands (Fig. SI 11).

Furthermore, it was investigated whether the ligands from the different MC runs bind to the protein in a similar fashion. To this end, the complexes of all the simulations were evaluated with LigPlot+ [46](SI Fig. SI 12). For the bromodomain protein the results show high similarity, all ligands form similar sets of hydrogen bonds with protein residues and similar contacts to surrounding protein chains. While no chemical similarity was found, there is some overlap in the way the ligands bind to the proteins, as shown in SI Fig. SI 13. To further investigated similarities in the distribution of known and generated ligands, UMAP[47] was employed through the ChemPlot implementation [48]. The approach allows for comparison of compounds in a principal component type of chemical space representation. As shown in figure SI 14 the distribution of the ligands is very diverse and, encouragingly, the ligands from the simulation are in all cases close to some clusters representing known ligands.

Finally, it was attempted to improve the SA, ESOL and QED score further, by adding a additional clean-up phase to the simulation. This consists of 100 simulation steps, with the score consisting only of the SA, ESOL and QED scores. However, under the condition that the Chai-1 score never fall below a certain threshold. The threshold was determined as the starting Chai-1 score minus 0.02. The probabilities for the types of steps were also adjusted. This clean up phase was tested for the best resulting structure found for each protein. The results are shown in SI 4. The SA, ESOL and QED score improve successfully, without incurring a major drawback in terms of the primary Chai-1 score. It is still important to note that even with these improvements, the SA score is only an estimate and does not guarantee that resulting structures can be synthesized with reasonable effort. Further work may be required to ensure the structures resulting from the simulation are reasonable.

### 3.4 Fragment based MC simulations

Instead of atomistic-step MC method the MC search based on larger chemical fragments was also investigated. In a first attempt some known ligands for the two proteins with high scores were selected. These ligands were broken down into fragments and recombined within a simulation of 500 steps, with *β* = 5. Hence, the expected outcome for this type of stochastic MC-based fragment assembly is to recover the known ligand (SI Table SI 5). Interestingly, for all cases compounds very similar to but not always exactly the same as the known ligands were recovered (SI Table SI 5). It reflects also the stochastic character of the compound generation.

Additionally, the MC simulation approach was tested on a set of 50 randomly selected fragments from the ChEMBL database [15] and fractured into BRICS fragments. These fragments were then used for a MC simulation of 1000 steps (SI (Table SI 6). Interestingly, the resulting scores are not as favorable as for the atomistic-step MC simulations. This is most likely due to the fact, that the fragments were chosen randomly and may not contain the most complementary chemical groups in a sterically well fitting fragment. In the future it might be possible to design smaller, more elementary fragments, that allow for a greater variety of combining chemical groups in different spatial arrangements.

However, it was also tested if fragment selections specifically suited for the selected target protein by decomposing known ligands result in improved performance. These fragments were then used in the same simulation setup as used for MC searches based on random fragments. Indeed, on average this type of MC simulation resulted in significantly improved scores compared to using randomly selected fragments (SI Table SI 7). The generated compounds indicate MMGBSA scores and Boltz-2 scores in the same range as known experimental binders to the target proteins. The course of the average MC confidence score and of the re-evaluation using MMGBSA was also investigated (SI Fig. SI 15). In both the MC simulations with random and specific fragments the scores improved rapidly at the beginning to reach MC scores around 0.7 to 0.8 and after 200 steps changed only gradually with significant fluctuations due to the relatively small *β* = 5 but also due the bigger changes caused by adding or removing whole fragments. The course of the MMGBSA score indicates similar trends (SI Fig. SI 15).

## 4 Conclusions

Recent advances in AI allow for the rapid and accurate structure prediction of proteins and also complexes with organic ligands[18, 21, 20]. It is also possible to use similar generative AI based methods to construct entirely new ligands employing, however, mostly rigid protein pockets as target input structures[7, 49, 3, 4]. Based on the rapid Chai-1 structure prediction approach we have developed an MC based approach, termed AI-MCLig, that searches stochastically through chemical space to generate new potential binders for a given target protein structure. The explicit rebuilding of the entire complex structure at every MC step allows for adaptation (induced fit) of the entire protein structure in response to an altered compound structure (and also of the ligand itself). This is a distinct advantage of our approach compared to existing construction methods that mostly construct ligands in a rigid protein environment[7].

For the targets in our study we found that only the average confidence score of Chai-1 correlates with experimentally measured ligand binding affinity. The limited accuracy of the score used to rapidly evaluate generated ligands clearly limits the applicability of the MC approach. However, this is an issue that affects all available docking and generative ligand design methods and we therefore consider this as a separate issue, not disfavoring our approach. This aspect clearly needs future efforts for improvement. Nevertheless, our approach allowed in all cases to rapidly suggest new potential binders with MMGBSA scores and Boltz-2 scores that are in the same range as those calculated for experimentally known ligands suggesting that these generated ligands potentially represent realistic binders.

We consider our MC based compound generation approach as complementary to the variety of fragment buildup and new reverse-diffusion based approaches. The method can be quickly adapted to specific needs. For example, one can limit the search just to modify side chains or subsets of positions of a known ligand and keep a desired core or reference structure unmodified. The method can be especially useful to rapidly generate a large variety of potential binders for a given target protein structure.

## Author contributions statement

J.A. performed simulations, wrote programs and scripts, analysed data, wrote paper draft. M.Z. designed project, supervised computational work, provided computational resources, wrote final manuscript version. J.A. and M.Z. analysed results, reviewed the manuscript.

## Acknowledgments

The authors thank Niklas Halbwedl and Patrick Quoika for useful discussions. This work was financially supported by the Deutsche Forschungsgemeinschaft (DFG Za153/29-1) and by local compute cluster (partially funded by DFG INST 95/1610-1 FUGG). We acknowledge additional HPC resources provided by the Erlangen National High Performance Center (NHR@FAU).

## References

[1] Javad Zahiria, Abbasali Emamjomehb, Samaneh Bagheri, et al. Protein complex prediction: A survey. Genomics, 112(1):174–183, 2020.

[2] Christopher Horvath. Comparison of preclinical development programs for small molecules (drugs/pharmaceuticals) and large molecules (biologics/biopharmaceuticals): studies, timing, materials, and costs. Pharmaceut Sci Encyclo: Drug Discovery, Development, and Manufacturing, pages 1–35, 2010.

[3] Pawewl ŚledŻ and Amedeo Caflisch. Protein structure-based drug design: from docking to molecular dynamics. Curr Opin Struct Biol, 48:93–102, 2018.

[4] Yidan Tang, Rocco Moretti, and Jens Meiler. Recent advances in automated structure-based de novo drug design. J Chem Inf Mod, 64(6):1794–1805, 2024.

[5] Elisa Martino, Sara Chiarugi, Francesco Margheriti, et al. Mapping, structure and modulation of ppi. Front Chem, 9, 2021.

[6] Mouchlis VD, Afantitis A, Serra A, et al. Advances in de novo drug design: From conventional to machine learning methods. Int J Mol Sci, 22(4)::1676, 2021.

[7] Xuhan Liu, Adriaan P IJzerman, and Gerard JP van Westen. Computational approaches for de novo drug design: past, present, and future. Artif Neu Networks, pages 139–165, 2020.

[8] Gabriel Corrëa Veríssimo, Rafaela Salgado Ferreira, and Vinícius Gonçalves Maltarollo. Ultralarge virtual screening: Definition, recent advances, and challenges in drug design. Mol Info, 44(1):e202400305, 2025.

[9] Oleksandr O Grygorenko, Dmytro S Radchenko, Igor Dziuba, et al. Generating multibillion chemical space of readily accessible screening compounds. Iscience, 23(11), 2020.

[10] Jiankun Lyu, John J Irwin, and Brian K Shoichet. Modeling the expansion of virtual screening libraries. Nature Chem Biol, 19(6):712–718, 2023.

[11] François Sindt, Guillaume Bret, and Didier Rognan. On the difficulty to rescore hits from ultralarge docking screens. J Chem Inf Mod, 2025.

[12] Petra Schneider, W Patrick Walters, Alleyn T Plowright, et al. Rethinking drug design in the artificial intelligence era. Nature Rev Drug Discov, 19(5):353–364, 2020.

[13] Christian Beier and Martin Zacharias. Tackling the challenges posed by target flexibility in drug design. Exp Opin Drug Discovery, 5(4):347–359, 2010.

[14] Xiangru Tang, Howard Dai, Elizabeth Knight, et al. A survey of generative ai for de novo drug design: new frontiers in molecule and protein generation. Brief Bioinfo, 25(4):bbae338, 2024.

[15] Barbara Zdrazil, Eloy Felix, Fiona Hunter, et al. The chembl database in 2023: a drug discovery platform spanning multiple bioactivity data types and time periods. Nucl Acids Res, 52(1):1180–1192, 11 2023.

[16] Zhihai Liu, Minyi Su, Liand Han, et al. Forging the basis for developing protein-ligand interaction scoring functions. Acc Chem Res, 50(2):302 – 309, 2017.

[17] Weixin Xie, Fanhao Wang, Yibo Li, et al. Advances and challenges in de novo drug design using three-dimensional deep generative models. J Chem Inf Mod, 62(10):2269–2279, 2022.

[18] John Jumper, Richard Evans, Alexander Pritzel, et al. Highly accurate protein structure prediction with alphafold. nature, 596(7873):583–589, 2021.

[19] Josh Abramson, Jonas Adler, Jack Dunger, et al. Accurate structure prediction of biomolecular interactions with alphafold 3. Nature, 630(8016):493–500, 2024.

[20] Jeremy Wohlwend, Gabriele Corso, Saro Passaro, et al. Boltz-1: Democratizing biomolecular interaction modeling. bioRxiv, pages 2024–11, 2024.

[21] Chai Discovery. Chai-1: Decoding the molecular interactions of life. bioRxiv, 2024.

[22] Jörg Degen, Christof Wegscheid-Gerlach, Andrea Zaliani, Matthias Rarey. On the art of compiling and using ‘drug-like’ chemical fragment spaces. ChemMedChem, 3(10):1503 – 1507, 2008.

[23] Greg Landrum. Rdkit: Open-source cheminformatics software. Zenodo, 2023.

[24] David Weininger. Smiles, a chemical language and information system. 1. introduction to methodology and encoding rules. J Chem Inf Comput Sci, 28(1):31–36, 1988.

[25] Peter Ertl, Ansgar Schuffenhauer. Estimation of synthetic accessibility score of drug-like molecules based on molecular complexity and fragment contributions. J Cheminfo, 1(8):302 – 309, 2009.

[26] John S. Delaney. Esol: Estimating aqueous solubility directly from molecular structure. J Chem Inf Comput Sci, 44(3):1000 – 1005, 2004.

[27] G.R. Bickerton, G.V. Paolini, J. Besnard, et al. Quantifying the chemical beauty of drugs. Nature Chem, 4:90–98, 2012.

[28] Bill R. III Miller, T. Dwight Jr. McGee, Jason M. Swails, et al. Mmpbsa.py: An efficient program for end-state free energy calculations. J Chem Theory Comput, 8(9):3314–3321, 2012.

[29] Wang, J., Wang, W., Kollman P.A.; Case, D.A. Automatic atom type and bond type perception in molecular mechanical calculations. J Mol Graph Mod, 25(2):247–60, 2006.

[30] Wang, J., Wolf, R. M.; Caldwell, J. W.; Kollman, P. A.; Case, D. A. Development and testing of a general amber force field. J Comput Chem, 25:1157–1174, 2006.

[31] David A. Case, Hasan Metin Aktulga, Kellon Belfon, et al. Amber 2025. University of California, San Francisco, 2025.

[32] David A. Case, Hasan Metin Aktulga, Kellon Belfon, et al. Ambertools. J Chem Inf Mod, 63(20):6183–6191, 2023. PMID: 37805934.

[33] William L Jorgensen, Jayaraman Chandrasekhar, Jeffry D Madura, et al. Comparison of simple potential functions for simulating liquid water. J Chem Phys, 79(2):926–935, 1983.

[34] Andreas W. Goetz, Mark J. Williamson Williamson, Dong Xu Xu, et al. Routine microsecond molecular dynamics simulations with AMBER - Part I: Generalized Born. J Chem Theo Comput, 8(5):1542–1555, 2012.

[35] Romelia Salomon-Ferrer Salomon-Ferrer, Andreas W. Goetz, Duncan Poole, et al. Routine microsecond molecular dynamics simulations with AMBER - Part II: Particle Mesh Ewald. J Chem Theo Comput, 9(9):3878–3888, 2013.

[36] Scott Le Grand; Andreas W. Goetz; Ross C. Walker. SPFP: Speed without compromise - a mixed precision model for GPU accelerated molecular dynamics simulations. Comp Phys Comm, 184:374–380, 2013.

[37] Richard Gowers, Max Linke, Jonathan Barnoud, et al. Mdanalysis: A python package for the rapid analysis of molecular dynamics simulations. In Proceedings of the 15th Python in Science Conference, SciPy, page 98–105. SciPy, 2016.

[38] Naveen Michaud-Agrawal, Elizabeth J. Denning, Thomas B. Woolf, et al. Mdanalysis: A toolkit for the analysis of molecular dynamics simulations. J Comput Chem, 32(10):2319–2327, April 2011.

[39] Douglas L. Theobald. Rapid calculation of rmsds using a quaternion-based characteristic polynomial. Acta Crystal, 61(4):478–480, June 2005.

[40] Pu Liu, Dimitris K. Agrafiotis, and Douglas L. Theobald. Fast determination of the optimal rotational matrix for macromolecular superpositions. J Compu Chem, 31(7):1561–1563, 2009.

[41] Stefan Van Der Walt, S Chris Colbert, and Gael Varoquaux. The numpy array: a structure for efficient numerical computation. Comput Sci & Eng, 13(2):22–30, 2011.

[42] Raymond E. Carhart, Dennis H. Smith, and R. Venkataraghavan. Atom pairs as molecular features in structure-activity studies: definition and applications. J Chem Info Comput Sci, 25(2):64–73, 1985.

[43] J. Eberhardt, D. Santos-Martins, A. F. Tillack, et al. Autodock vina 1.2.0: New docking methods, expanded force field, and python bindings. J Chem Inf Mod, 61(8):3891–3898, 8 2021.

[44] A. J. Olson O. Trott. Autodock vina: improving the speed and accuracy of docking with a new scoring function, efficient optimization, and multithreading. J Comput Chem, 31(2):455–661, 1 2010.

[45] Saro Passaro, Gabriele Corso, Jeremy Wohlwend, et al. Boltz-2: Towards accurate and efficient binding affinity prediction. 2025.

[46] Roman A. Laskowski and Mark B. Swindells. Ligplot+: Multiple ligand–protein interaction diagrams for drug discovery. J Chem Inf Mod, 51(10):2778–2786, 2011.

[47] L. McInnes, J. Healy, and J. Melville. UMAP: Uniform Manifold Approximation and Projection for Dimension Reduction. ArXiv e-prints, February 2018.

[48] Murat Cihan Sorkun, Dajt Mullaj, J. M. Vianney A. Koelman, and Süleyman Er. Chemplot, a python library for chemical space visualization. Chemistry–Methods, 2(7):e202200005, 2022.

[49] Mingyang Wang, Zhe Wang, Huiyong Sun, et al. Deep learning approaches for de novo drug design: An overview. Curr Opin Struct Biol, 72:135–144, 2022.

