## Supplementary Material for "De novo protein ligand design including protein flexibility and conformational adaptation"

Jakob Agamia[1] and Martin Zacharias[1]
[1]Center of Functional Protein Assemblies and Physics Department, Technical University Munich, Ernst-Otto-Fischer-Str. 8, 85748, Bavaria, Germany

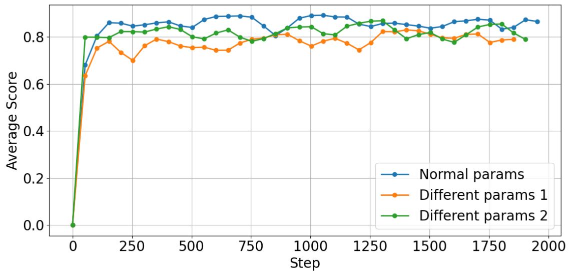


Figure SI 1: The development of the score during a atomistic-step MC simulation on the bromodomain protein for some different choices of parameters for likelihood of each type of steps. As shown the standard parameters outperform the other choices. The probabilities for the first set of different parameters were: Adding side groups=0.2, Removing atoms=0.35, Changing atom types=0.1, Adding atoms in chain=0.1, Changing bond types=0.025, Forming rings=0.025, Breaking up rings=0.05, Turning rings aromatic=0.05, Rearranging bonds=0.1. And for the second set of different parameters they were: Adding side groups=0.2, Removing atoms=0.2, Changing atom types=0.15, Adding atoms in chain=0.1, Changing bond types=0.05, Forming rings=0.05, Breaking up rings=0.05, Turning rings aromatic=0.1, Rearranging bonds=0.1.

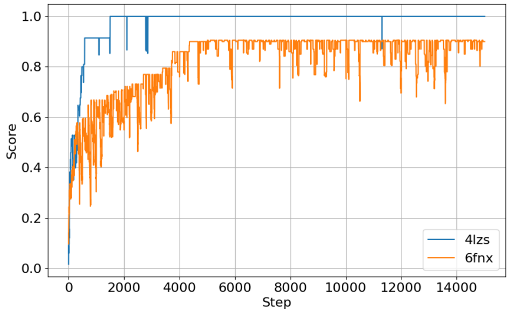


Figure SI 2: Generation of target compounds during atomistic-step MC searches starting from benzene and using the dice similarity score (y-axis). The target example compounds were taken from pdb4lzs (blue) and pdb6fnx (orange) and a benzene molecule was used as start structure. Each MC step (x-axis) consists of chemical changes in the compound (see Methods) that are accepted or rejected according to the dice similarity with respect to the target compound. A dice score of 1 indicates chemical identity.

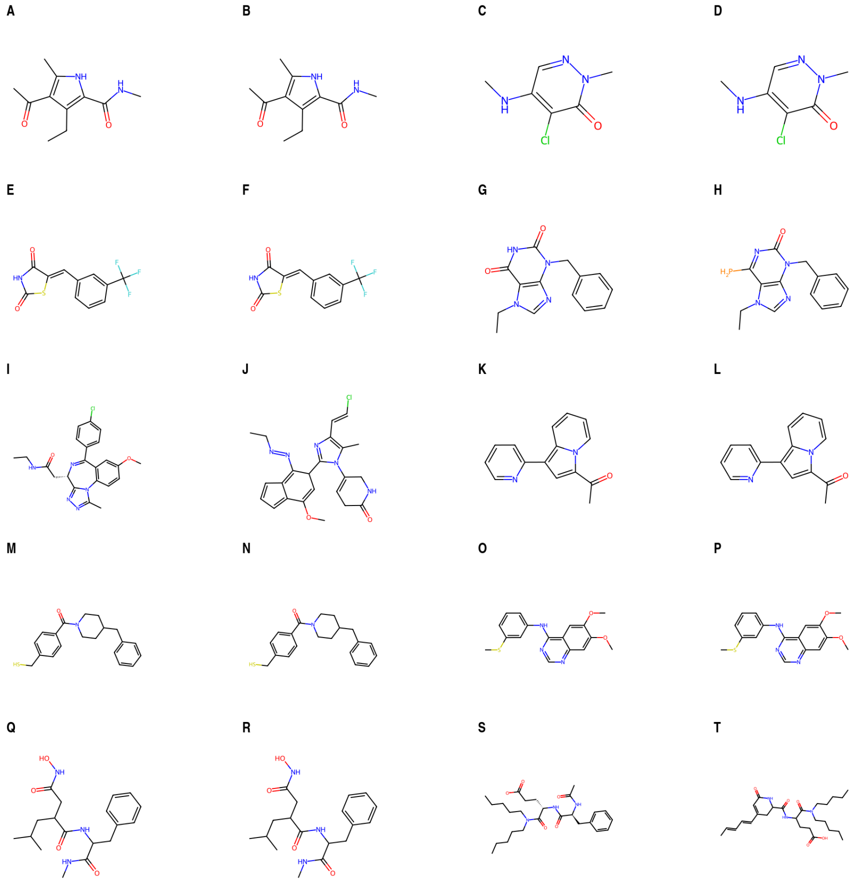


Figure SI 3: Chemical structures of the target ligands and reconstructed ligands starting from benzene for the MC simulation based on atom and bond type changes. Target (A) and reconstructed (B) ligand for the compound in pdb4lzs complex; (C, D) same for the pdb5mli complex; (E, F) same for the pdb3vc4 complex; (G, H) same for the pdb6fnx complex; (I, J) same for the pdb2yek complex; (K, L) same for the pdb4a9i complex; (M, N) same for the pdb3iw7 complex; (O, P) same for the pdb1di9 complex; (Q, R) same for the pdb1mnc complex; (S, T) same for the pdb1a07 complex.

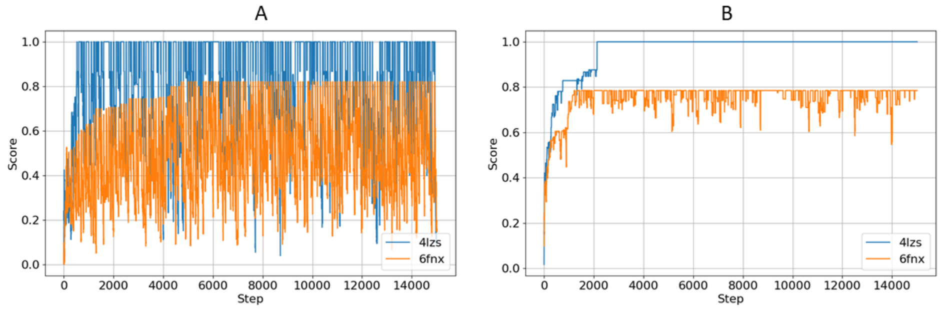


Figure SI 4: Test of different simulation temperature parameters $\beta$ for the recovery of chemical compounds based on the atomistic-step MC search using a similarity dice score as target score. Results are shown for target compounds extracted from pdb4lzs (blue) and pdb 6fnx (orange), (A) MC simulation with $\beta = 20$, (B) MC simulation with $\beta = 70$. The smaller $\beta = 20$ results in large fluctuations of the score compared to $\beta = 70$. Overall both conditions perform worse than the optimal choice of $\beta = 50$, see main text.

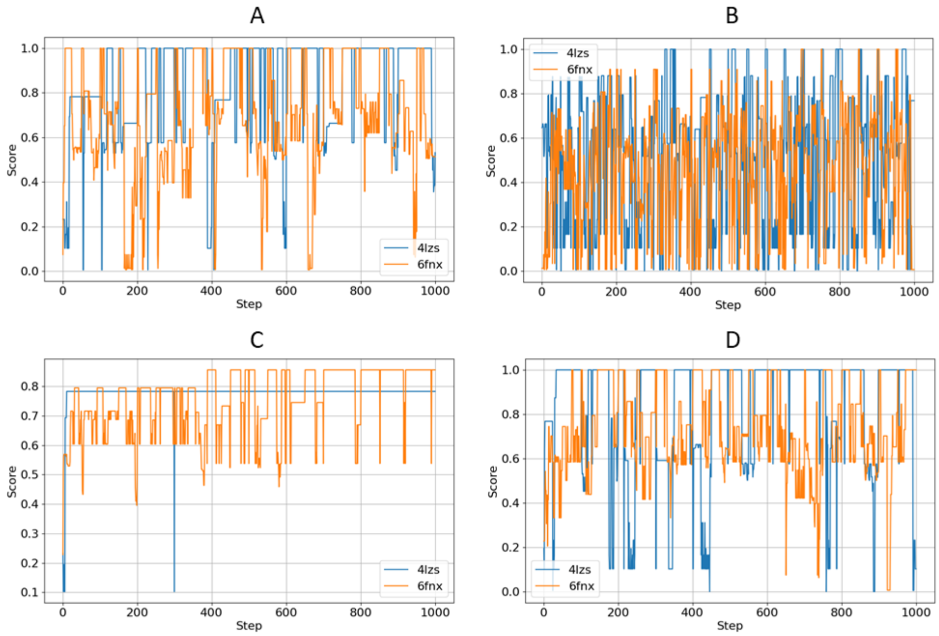


Figure SI 5: Dice similarity score during the fragment based simulation for the compounds extracted from pdb4lzs (blue) and pdb6fnx (orange), for (A) a $\beta = 5$, (B)$\beta = 2$, (C) $\beta = 10$, (D) $\beta = 5$ but using a probability of 0.7 for adding fragments and 0.3 for removing fragments.

Table SI 1: Targeted and resulting compounds in SMILES representation for the atomistic-step MC simulations for a variation of the MC probability parameters: Adding side groups=0.2, Removing atoms=0.35, Changing atom types=0.1, Adding atoms in chain=0.1, Changing bond types=0.025, Forming rings=0.025, Breaking up rings=0.05, Turning rings aromatic=0.05, Rearranging bonds=0.1. MC simulations were started from benzene and the dice similarity score was used. The target compound were extracted from the pdb-entries given in column 1 and were taken from [16]. Simulation setup was the same as for the standard MC simulation parameters as explained in the main text

| Complex | Target | Result | Dice score |
|---|---|---|---|
| 4lzs | CNC(=O)c1[nH]c(c(c1CC)C(=O)C)C | CCc1c(C(=O)NC)[nH]c(C)c1C(C)=O | 1.00 |
| 5mli | CNc1cnn(c(=O)c1Cl)C | CNc1c(Cl)c(=O)cnn1C | 0.836 |
| 3vc4 | O=C1NC(=O)/C(=C/c2cccc(c2)C(F)(F)F)/S1 | O=C1C=C(C=C2C=C3[PH](=C2)C=CC3(F)F)C(=O)N1 | 0.765 |
| 6fnx | CCn1cnc2c1c(=O)[nH]c(=O)n2Cc1ccccc1 | CCn1cnc2c1c(=O)nc(N)n2Cc1ccccc1 | 0.905 |
| 2yek | CCNC(=O)C[C@@H]1N=C(c2ccc(cc2)Cl)c2c(-n3c1nnc3C)ccc(c2)OC | CCNn1cnc2c1=C(C1=NCC3=CC(OC)=CC3=C1C1=C[C+]=C(Cl)C=C1)C(C(C)=O)N=2 | 0.748 |
| 4a9i | CC(=O)c1cc(c2n1cccc2)c1ccccn1 | CC(=O)c1cccc[n+]1-c1cc[nH]c1C=CC=O | 0.850 |
| 3iw7 | SCc1ccc(cc1)C(=O)N1CCC(CC1)Cc1ccccc1 | O=C(c1ccc(CS)cc1)N1CCC(Cc2ccccc2)CC1 | 1.00 |
| 1di9 | COc1cc2c(ncnc2cc1OC)Nc1cccc(c1)SC | COc1[nH]cc(-c2cc[nH]c2Nc2cccc(SC)c2)c1OC | 0.909 |

Table SI 2: Targeted and resulting compounds in SMILES representation for the atomistic-step MC simulations for a variation of the MC probability parameters: Adding side groups=0.2, Removing atoms=0.2, Changing atom types=0.15, Adding atoms in chain=0.1, Changing bond types=0.05, Forming rings=0.05, Breaking up rings=0.05, Turning rings aromatic=0.1, Rearranging bonds=0.1. MC simulations were started from benzene and the dice similarity score was used. The target compound were extracted from the pdb-entries given in column 1 and were taken from [16]. Simulation setup was the same as for the standard MC simulation parameters as explained in the main text

| Complex | Target | Result | Dice score |
|---|---|---|---|
| 4lzs | CNC(=O)c1[nH]c(c(c1CC)C(=O)C)C | CCc1c(C(=O)NC)[nH]c(C)c1C(C)=O | 1.00 |
| 5mli | CNc1cnn(c(=O)c1Cl)C | CNc1cnn(C)c(=O)c1Cl | 1.00 |
| 3vc4 | O=C1NC(=O)/C(=C/c2cccc(c2)C(F)(F)F)/S1 | O=CC1=CC(=CC2=C(F)C(=O)N[SH2]2)C=CC1(F)F | 0.797 |
| 6fnx | CCn1cnc2c1c(=O)[nH]c(=O)n2Cc1ccccc1 | CC[N+]1=c2c(n(Cc3ccccc3)c3nncn23)=NC1=O | 0.800 |
| 2yek | CCNC(=O)C[C@@H]1N=C(c2ccc(cc2)Cl)c2c(-n3c1nnc3C)ccc(c2)OC | CCNC1=CC(C2=C3C(C=COC)=CC=c4[nH]c(Cl)cc4=C3OCC(C)=N2)N=C1 | 0.728 |
| 4a9i | CC(=O)c1cc(c2n1cccc2)c1ccccn1 | CC(=O)c1cc(-c2ccccc2)c2[nH]ccn12 | 0.900 |
| 3iw7 | SCc1ccc(cc1)C(=O)N1CCC(CC1)Cc1ccccc1 | O=C(c1ccc(CS)cc1)N1CCC(Cc2ccccc2)CC1 | 1.00 |
| 1di9 | COc1cc2c(ncnc2cc1OC)Nc1cccc(c1)SC | COc1ccc(C2=CN=CN=C3C=C(SC)C=C3N2)cc1OC | 0.877 |

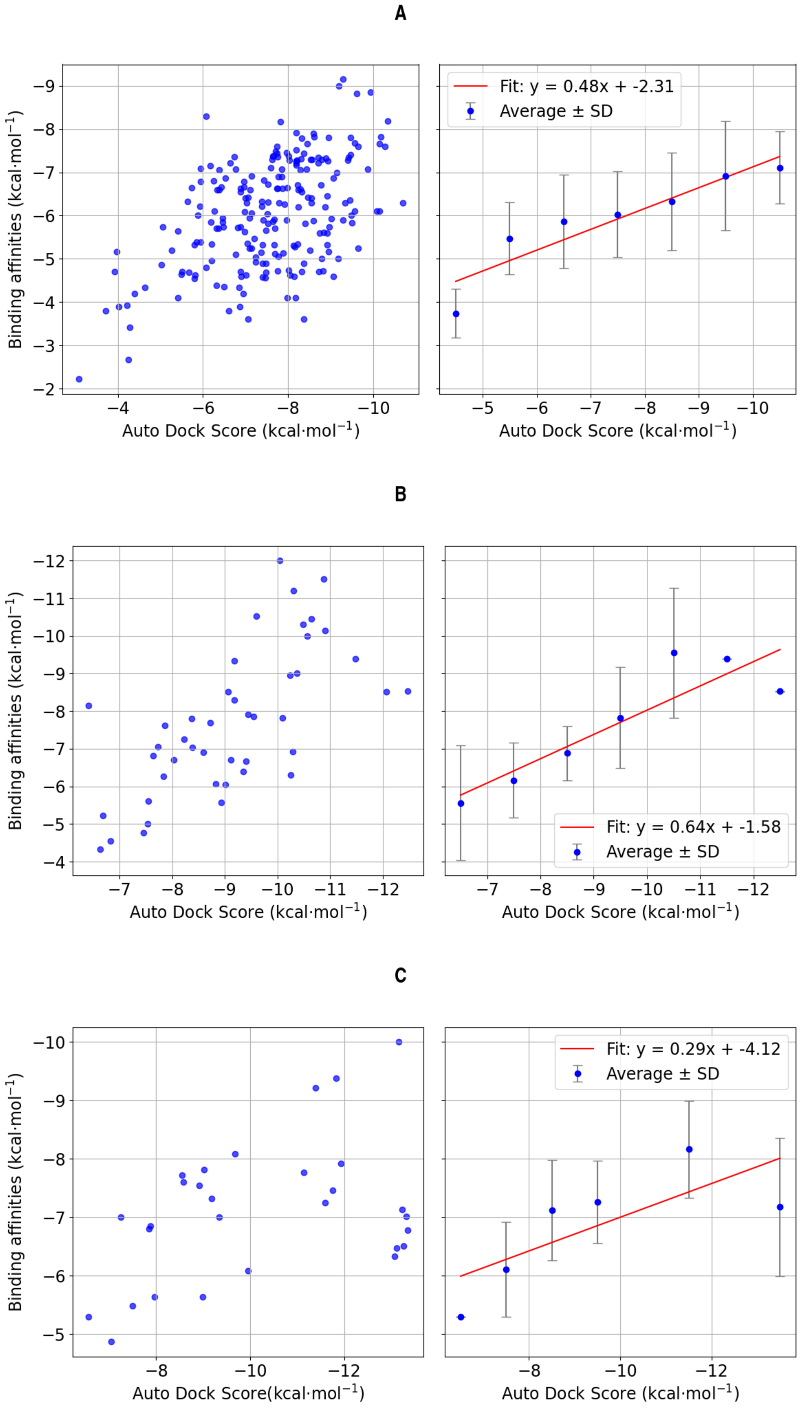


Figure SI 6: Correlation of the experimentally determined compound binding affinity (extracted from [16]) and the Auto-dock score[43]. (A) (left panel) Data for different ligands bound to the bromodomain, (right panel) averages for splitting the data into 8 score segments (B) same as (A) for ligands of the serine/threonine-protein kinase pim-1, (C) same as (A) for ligands of the p38 map kinase complex.

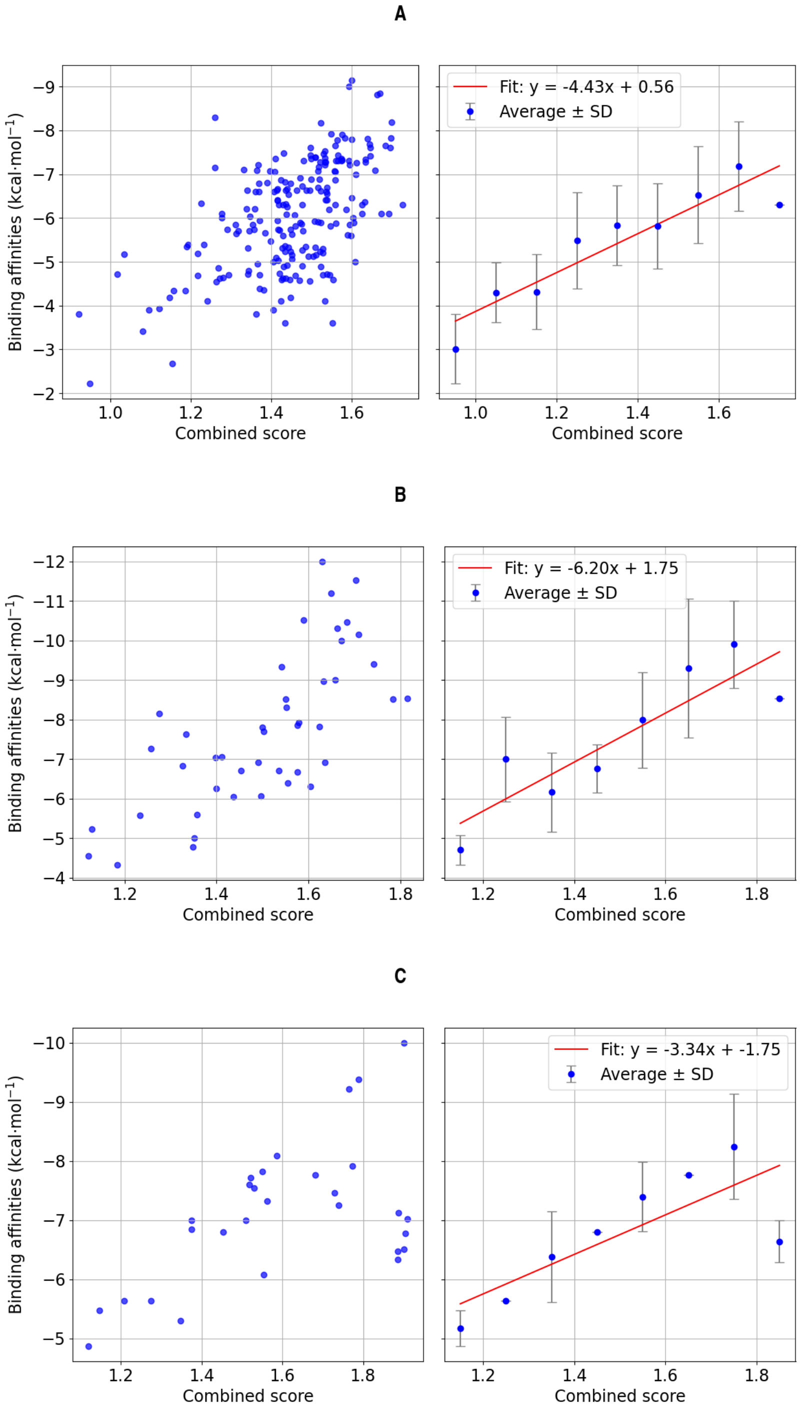


Figure SI 7: Correlation of the experimentally determined compound binding affinity (extracted from [16]) and the sum of Chai-1 confidence and normalized Auto-dock score[43]. (A) (left panel) Data for different ligands bound to the bromodomain, (right panel) averages for splitting the data into 8 score segments (B) same as (A) for ligands of the serine/threonine-protein kinase pim-1, (C) same as (A) for ligands of the p38 map kinase complex. (C) Binding affinity against the combined score for ligands of the p38 map kinase complex.

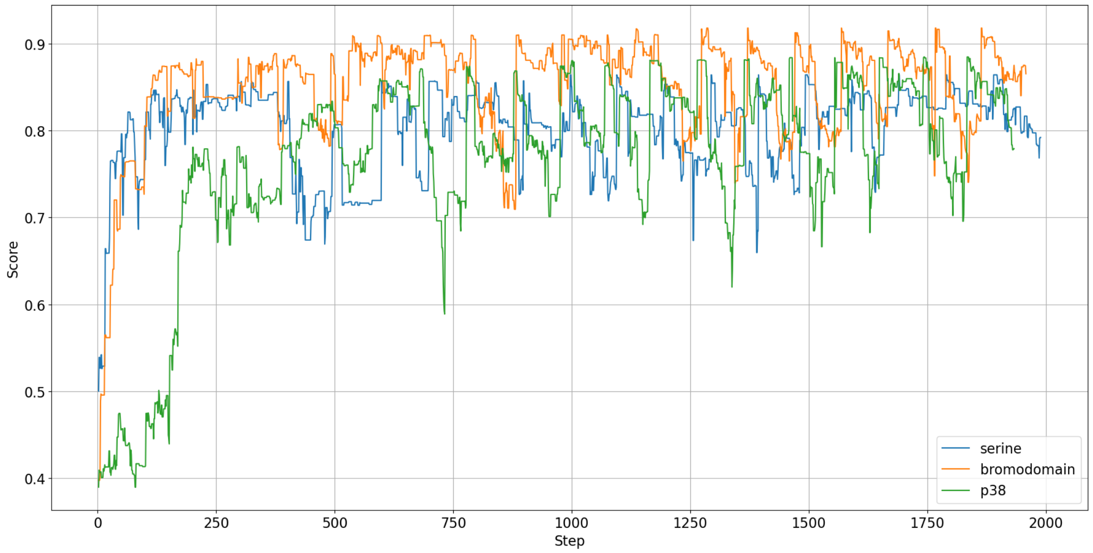


Figure SI 8: The evolution of the confidence score during the atomistic-step MC simulation for one of the three MC runs for each target protein.

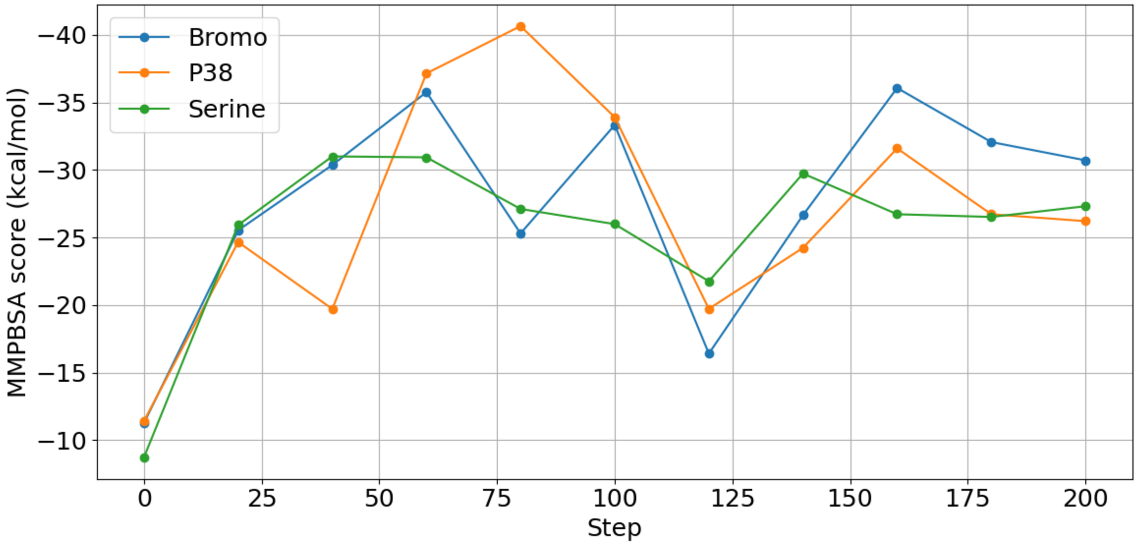


Figure SI 9: Re-evaluation of generated compounds during first 200 MC steps using the MMGBSA score (indicated for one trial run for each target protein

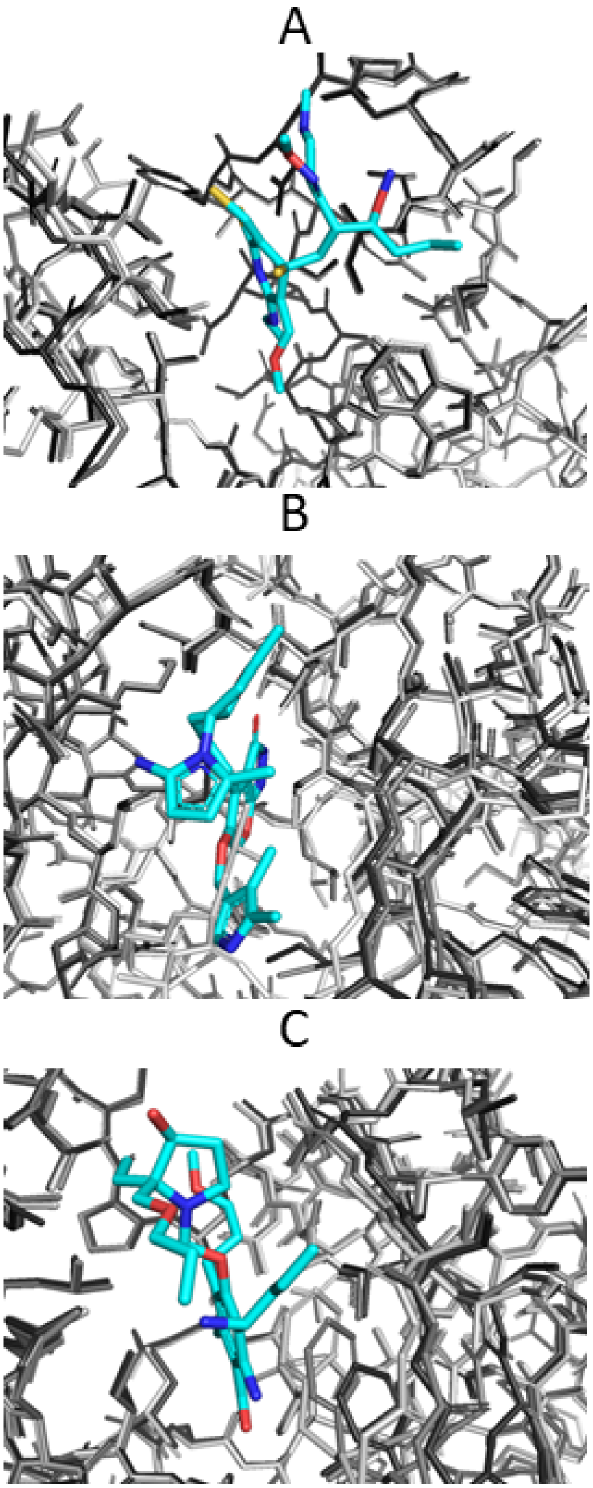


Figure SI 10: Illustration of conformational protein structure adjustment during MC based ligand generation. (A) Best superposition of the bromodomain protein ligand binding pocket after zero (light grey stick model), 100 (grey sticks) and 2000 (dark grey sticks) MC steps. For clarity only the final generated ligand is shown as blue stick model. (B) same as (A) but for the p38 map kinase target. (C) same as (A) but for the serine/threonine pim-1 kinase target.

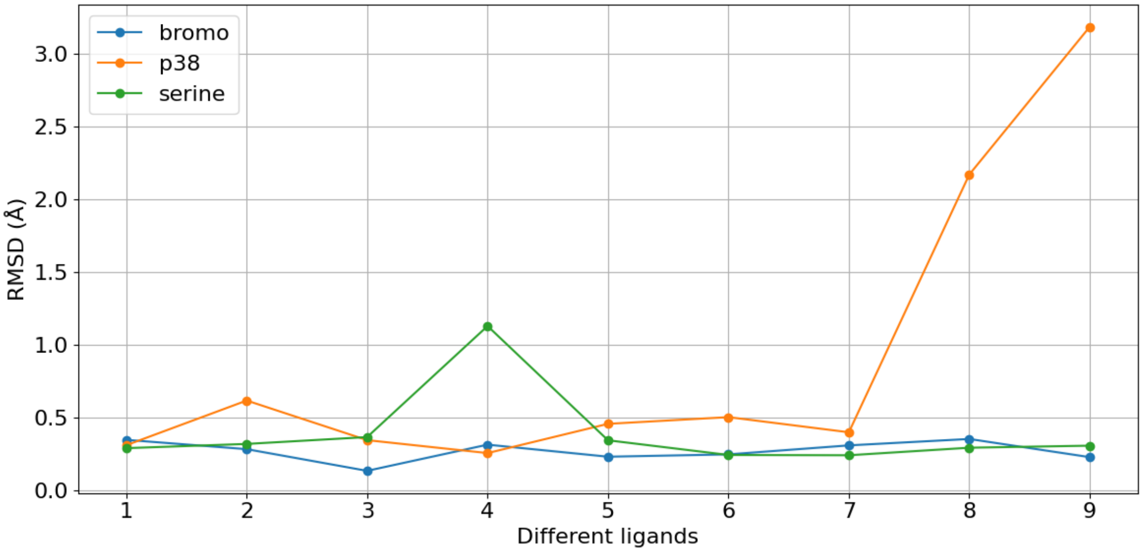


Figure SI 11: The RMSD of the protein binding pocket for some experimentally known ligand-protein complexes for each of the target proteins (the pocket atoms were defined as all atoms within 5 of the ligand).

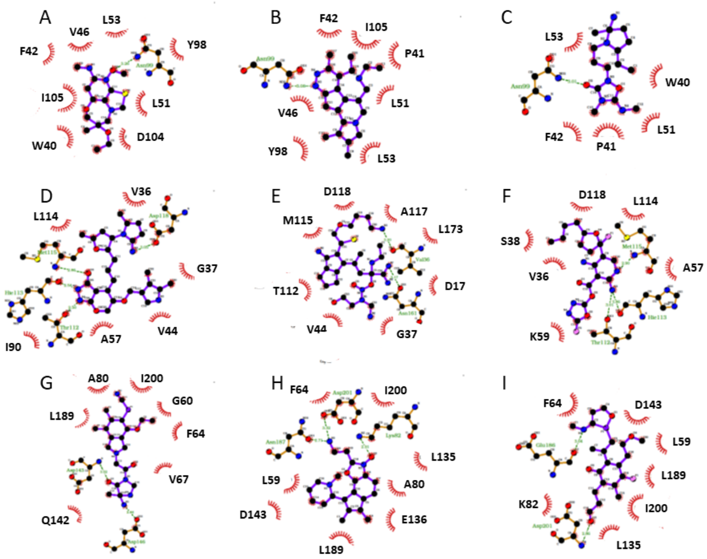


Figure SI 12: LigPlot[46] representation of the final generated ligands. (A-C) Results for the 3 MC runs for the bromodomain case (D-F) Final generated ligands for the p38 map kinase case. (G-H) same for Pim-1 kinase target.

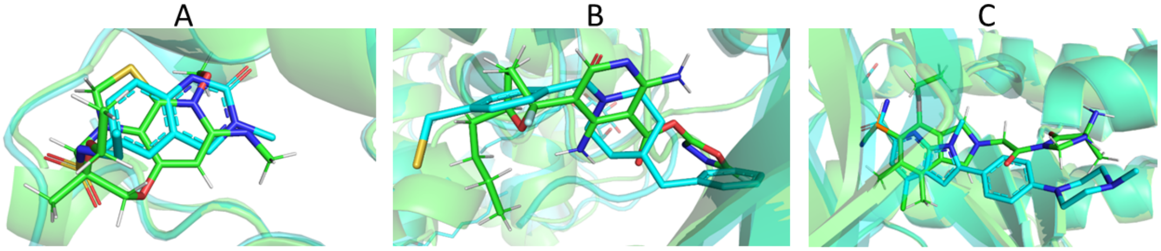


Figure SI 13: Examples of generated structures compared with known binders. (A) Final designed ligand from first bromodomain MC run (green sticks) and similar experimental binder (blue sticks). (B) same as (A) for the p38 kinase case. (C) same as (A) but for Pim-1 kinase case.

Table SI 3: The target and resulting compounds in SMILES representation for the fragment-based MC simulations. A random fragment of the target ligand was used as starting compound and the dice similarity score was optimized in the MC search. The target SMILES correspond to bromodomain ligands extract from the PDB-bind [16].

| **Target(Smiles) (from pdb)** | **Final closest compound** | **Dice score** |
|---|---|---|
| `CNC(=O)c1[nH]c(c(c1CC)C(=O)C)C` (4lzs) | `CCc1c(C(=O)NC)[nH]c(C)c1C(C)=O` | 1.00 |
| `CNc1cnn(c(=O)c1Cl)C` (5mli) | `CNc1cnn(C)c(=O)c1Cl` | 1.00 |
| `O=C1NC(=O)/C(=C/c2cccc(c2)C(F)(F)F)/S1` (3vc4) | `O=C1NC(=O)C(=Cc2cccc(C(F)(F)F)c2)S1` | 1.00 |
| `CCn1cnc2c1c(=O)[nH]c(=O)n2Cc1ccccc1` (6fnx) | `CCn1cnc2c1c(=O)[nH]c(=O)n2Cc1ccccc1` | 1.00 |
| `CCNC(=O)C[C@@H]1N=C(c2ccc(cc2)Cl)c2c(-n3c1nnc3C)ccc(c2)OC` (2yek) | `CCNC(=O)C[C@@H]1N=C(c2ccc(Cl)cc2)c2cc(OC)ccc2-n2c(C)nnc21` | 1.00 |
| `CC(=O)c1cc(c2n1cccc2)c1ccccn1` (4a9i) | `CC(=O)c1cc(-c2ccccn2)c2ccccn12` | 1.00 |
| `SCc1ccc(cc1)C(=O)N1CCC(CC1)Cc1ccccc1` (3iw7) | `O=C(c1ccc(CS)cc1)N1CCC(Cc2ccccc2)CC1` | 1.00 |
| `COc1cc2c(ncnc2cc1OC)Nc1cccc(c1)SC` (1di9) | `COc1cc2ncnc(Nc3cccc(SC)c3)c2cc1OC` | 1.00 |
| `CNC(=O)[C@H](Cc1ccccc1)NC(=O)[C@@H](CC(=O)NO)CC(C)C` (1mnc) | `CNC(=O)[C@@H](Cc1ccccc1)NC(=O)[C@@H](CC(=O)NO)CC(C)C` | 1.00 |
| `CCCCCN(C(=O)[C@@H](NC(=O)[C@H](Cc1ccccc1)NC(=O)C)CCC(=O)[O-])CCCCC` (1a07) | `N(C(C)=O)[C@H](Cc1ccccc1)C(=O)N[C@H](CCC(=O)[O-])C(=O)N(CCCCC)CCCCC` | 1.00 |

Table SI 4: Final generated compounds (Smiles code, some of the Smiles are split into two lines) after the clean up phase, for the bromodomain, p38 kinase and serine/threonine-protein kinase pim-1 and the final scores.

| **Attempt** | **Results for bromodomain** | **Score** | **Chai-1 score** | **SA score** | **ESOL score** | **QED score** |
|---|---|---|---|---|---|---|
| 2 | `CCOc1c(C)nnc2c1C(CC(C)N1CCC(O)CC1)N(C)C2` | 0.909 | 0.900 | 3.926 | -2.969 | 0.887 |
| **Attempt** | **Results for p38 map kinase** | | | | | |
| 1 | `CCc1c(C2CCN(CC)CC2)noc2cnc(O)c1-2` | 0.878 | 0.861 | 2.950 | -3.346 | 0.933 |
| **Attempt** | **Results for Pim-1 kinase** | | | | | |
| 2 | `CCn1c(C)c2ccc3c(c2c1N1CCOC1)CN(C)N3CO` | 0.865 | 0.843 | 3.687 | -3.078 | 0.937 |
| **Attempt** | **Results for $\beta$-1 receptor** | | | | | |
| 3 | `Cc1c2ccc(CN3CCCC3)cc2c(CC(=O)CO)n1C` | 0.869 | 0.849 | 2.715 | -3.053 | 0.920 |

Table SI 5: The ligands resulting from the fragment simulation for single fragmented ligands

| **Complex** | **Fragmented ligand** | **Result** | **Score** | **Dice score** |
|---|---|---|---|---|
| 2yek/bromodomain | `CCNC(=O)C[C@@H]1N=C(c2ccc(cc2)Cl)c2c(-n3c1nnc3C)ccc(c2)OC` | `[5*]NC(=O)C[C@@H]1N=C(c2ccc(Cl)cc2)c2cc(OC)ccc2-n2c(C)nnc21` | 0.919 | 0.895 |
| 5cqt/bromodomain | `CCN1c2ccc(c3c2c(C1=O)ccc3)S(=O)(=O)NC1CCCCC1` | `CCN1C(=O)c2cccc3c(S(=O)(=O)NC4CCCCC4)ccc1c23` | 0.9245 | 1.0 |
| 6fnx/bromodomain | `CCn1cnc2c1c(=O)[nH]c(=O)n2Cc1ccccc1` | `[9*]n1cnc2c1c(=O)[nH]c(=O)n2Cc1ccccc1` | 0.916 | 0.695 |
| 3gc7/p38 kinase | `Fc1ccc(c(c1)Cl)c1cc(cc2c1cnc(=O)n2c1c(Cl)cccc1Cl)[C@@H]1CC[N@H+](CC1)C(C)C` | `[5*][N@H+]1CC[C@@H](c2cc(-c3ccc(F)cc3Cl)c3cnc(=O)n([C@H]4CC[N@@H+]([5*])CC4)c3c2)CC1` | 0.896 | 0.619 |
| 6ohd/p38 kinase | `Cc1ccc(cc1c1cnc2c(c1)[nH]c(=O)n2C(C)(C)C)C(=O)Nc1nocc1` | `Cc1ccc(-c2ccon2)cc1-c1cnc2c(c1)[nH]c(=O)n2C(C)(C)C` | 0.862 | 0.690 |
| 3hv5/p38 kinase | `O=C(Nc1cc(nn1c1ccc(cc1)C)C(C)(C)C)Nc1cccc(c1)Nc1cc[nH+]c2c1cc(cc2)[N+](=O)[O-]` | `[16*]c1cccc(-c2cc(C(C)(C)C)nn2-c2cc[nH+]c3ccc([N+](=O)[O-])cc23)c1` | 0.846 | 0.459 |
| 4dtk/Pim-1 kinase | `CC(Oc1cccc(c1N1CCC[C@H](C1)[NH3+])/C=C/1\SC(=O)NC1=O)C` | `[16*]c1cccc([16*])c1-c1cccc(C=C2SC(=O)NC2=O)c1N1CCC[C@@H]([NH3+])C1` | 0.865 | 0.684 |
| 4n6z/Pim-1 kinase | `O=C(c1nccnc1N)Nc1c[nH+]ccc1N1CCC[C@@H](C1)[NH3+]` | `Nc1nccnc1C(=O)Nc1c[nH+]ccc1N1CCC[C@H]([NH3+])C1` | 0.870 | 1.0 |
| 5kzi-1 kinase | `Fc1cccc(c1c1ccc2n(n1)c(nc2)Nc1c[nH+]ccc1N1CCC[C@@H](C1)[NH3+])F` | `[NH3+][C@H]1CCCN(c2cc[nH+]cc2-c2ccc3cnc(Nc4c(F)cccc4F)n3n2)C1` | 0.851 | 0.649 |

A

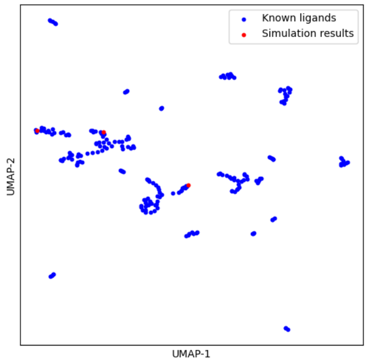


B

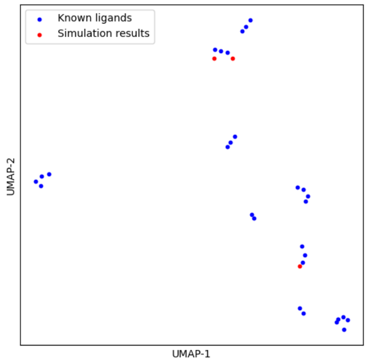


C

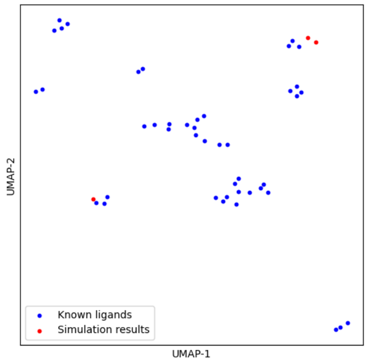


Figure SI 14: UMAP analysis of the chemical space for the known ligand of each protein and the results from the atom-based simulation.

Table SI 6: The ligands resulting from the fragment based simulation, with random fragments, and the final scores

| Attempt | Results for bromodomain | Score | Chai-1 score | SA score | ESOL score | QED score | MMPBSA score $\frac{kcal}{mol}$ | Boltz-2 score $\frac{kcal}{mol}$ |
|---|---|---|---|---|---|---|---|---|
| 1 | [5*]N1CCN(c2ccc3ncn(C[8*])c(=O)c3c2)CC1 | 0.907 | 0.899 | 2.250 | -1.916 | 0.786 | $-28.13 \pm 0.11$ | -8.157 |
| 2 | [8*]CCCN1CCC1(C)N1CCC(c2n[nH]c(=O)c3ccccc23)CC1 | 0.884 | 0.870 | 3.454 | -3.831 | 0.930 | $-29.15 \pm 0.11$ | -8.393 |
| 3 | [7*]C[C@H]1[C@H](O)[C@H](CCCN2CCC[C@H]([15*])C2)[C@H] | | | | | | | |
| (OC(F)(F)F)O[C@H]1c1cc2[nH]cc([16*])c(=O)c2cc1N | 0.900 | 0.910 | 4.894 | -5.008 | 0.543 | $-28.20 \pm 0.12$ | -8.294 | |
| **Attempt** | **Results for p38 map kinase** | | | | | | | |
| 1 | [15*]C1CCN(C2=Nc3cc(-c4cc([16*]cc(NC=O)c4)c([16*])cc3) | | | | | | | |
| S(C)(=O)=N2)CC1 | 0.836 | 0.820 | 3.485 | -4.778 | 0.819 | $-28.55 \pm 0.12$ | -7.827 | |
| 2 | Cn1c(=O)c(N2CCCCC2)cc2cnc(-c3cccc(B(O)O)c3)nc21 | 0.863 | 0.854 | 2.790 | -2.758 | 0.664 | $-27.71 \pm 0.15$ | -8.453 |
| 3 | [5*]N([5*])C(C)c1cc(C2C(C(=O)N3CCOCC3c3ccccc3C) | | | | | | | |
| (=O)N2c2ncnc(N)c2CN)c(N(C)C)s1 | 0.855 | 0.853 | 4.313 | -4.446 | 0.418 | $-28.40 \pm 0.12$ | $-7.884$ | |
| **Attempt** | **Results for Pim-1 kinase** | | | | | | | |
| 1 | [5*]NC(=O)c1cc(-c2ccccc2[16*])cc([C@H] | | | | | | | |
| 2CCN([5*])C2)c1[16*] | 0.846 | 0.824 | 2.543 | -3.330 | 0.897 | $-30.68 \pm 0.12$ | -9.046 | |
| 2 | [5*]N(C)S(=O)(=O)Cc1cncc(N2Cc3ccccc3C2=O)c1 | 0.853 | 0.827 | 2.501 | -2.761 | 0.921 | $-30.08 \pm 0.12$ | -9.067 |
| 3 | [5*]N1CCOC[C@H](N2CC3CN(c4ccc5c(c4)cc4n5CCC4CC(=O)O)CC3 | | | | | | | |
| C2)C1 | 0.862 | 0.841 | 4.213 | -3.658 | 0.782 | $-32.27 \pm 0.12$ | -8.135 | |
| **Attempt** | **Results for $\beta$-1 receptor** | | | | | | | |
| 1 | [5*]N1CCCc2c1sc(-c1ccc(=O)n(C)c1)c2CN | 0.834 | 0.812 | 3.145 | -3.361 | 0.866 | $-27.42 \pm 0.15$ | -6.806 |
| 2 | [[14*]c1nnc(N2CC(O)CN(c3cc(O)nc(-c4c(Cl) | | | | | | | |
| cc(Cl)c5ccc(C)nc45)n3)CC2=O)s1 | 0.842 | 0.845 | 3.844 | -5.252 | 0.401 | $-46.01 \pm 0.22$ | -8.696 | |
| 3 | [1*]C(=O)C1c2ccccc2C(=O)N(c2cc(C[C@H](N)C[4*])ccN3Cc4ccc | | | | | | | |
| cc4N(([5*])[C@@H]([13*])[C@@H]3C)n2)C1[13*] | 0.843 | 0.841 | 4.315 | -5.251 | 0.552 | $-38.06 \pm 0.19$ | -9.835 | |

Table SI 7: The ligands resulting from the fragment based simulation, with fragments from known ligands and the final scores

| Attempt | Results for bromodomain | Score | Chai-1 score | SA score | ESOL score | QED score | MMPBSA score $\frac{kcal}{mol}$ | Boltz-2 score $\frac{kcal}{mol}$ |
|---|---|---|---|---|---|---|---|---|
| 1 | CCN1C(=O)c2cccc3c([NH+]4CCCC4)ccc1c23 | 0.915 | 0.910 | 3.656 | -3.078 | 0.886 | $-29.70 \pm 0.12$ | -8.720 |
| 2 | Cc1nnc2n1-c1ccc(N3CC[NH2+]CC3)cc1C(c1ccc(Cl)cc1) | | | | | | | |
| =N[C@H]2[C@@H](C)C(=O)[NH+]1CCCCC1 | 0.910 | 0.911 | 4.874 | -4.312 | 0.535 | $-28.22 \pm 0.11$ | -8.835 | |
| 3 | CCC(=O)N[C@@H]1N=C(c2ccccc2)c2ccccc2-n2c(C)nnc21 | 0.913 | 0.918 | 2.872 | -4.127 | 0.793 | $-28.01 \pm 0.12$ | -9.376 |
| **Attempt** | **Results for p38 map kinase** | | | | | | | |
| 1 | [9*]n1c(=O)[nH]c2cc(-c3cc(C(N)=O)ccc3Cl)cnc21 | 0.879 | 0.874 | 2.978 | -3.045 | 0.751 | $-29.93 \pm 0.17$ | -7.733 |
| 2 | [8*]C[N@H+]1CC[C@@H](c2nc(-c3c(F)cc(Br)cc3F)c | | | | | | | |
| (-c3ccncc3)[nH]2)CC1 | 0.876 | 0.876 | 3.590 | -5.051 | 0.662 | $-41.96 \pm 0.14$ | -11.107 | |
| 3 | [16*]c1ccc(F)c(-c2nc(-n3nc(N4CCOCC4)cc3C3CC[NH2+] | | | | | | | |
| CC3)[nH]c2-c2ccc3c(c2)OCCO3)c1 | 0.860 | 0.866 | 3.610 | -5.187 | 0.411 | $-38.67 \pm 0.18$ | -8.878 | |
| **Attempt** | **Results for Pim-1 kinase** | | | | | | | |
| 1 | Cn1c(C=C2SC(NC[C@H]3C[NH2+]CCC34CC4) | | | | | | | |
| =NC2=O)cc2c(C#N)ccnc21 | 0.860 | 0.842 | 4.975 | -3.260 | 0.749 | $-44.70 \pm 0.11$ | -10.911 | |
| 2 | [14*]c1nc(-c2cc(O)n3ncc(N4CC[NH+](C)CC4)c3n2)ccc1F | 0.869 | 0.859 | 3.825 | -2.194 | 0.691 | $-29.11 \pm 0.11$ | -8.702 |
| 3 | [14*]c1nc(-c2cncc([C@@H]3CCCC[C@H]3[NH3+])n2)ccc1F | 0.868 | 0.849 | 3.877 | -3.073 | 0.909 | $-21.06 \pm 0.13$ | -8.153 |
| **Attempt** | **Results for $\beta$-1 receptor** | | | | | | | |
| 1 | O=c1[nH]c2cccc([C@H](O)CN3CCOCC3)c2[nH]1 | 0.843 | 0.822 | 2.973 | -1.765 | 0.740 | $-21.81 \pm 0.16$ | -8.038 |
| 2 | O=c1[nH]c2cccc(N3CC[NH2+]CC3)c2[nH]1 | 0.853 | 0.840 | 3.725 | -1.071 | 0.590 | $-33.00 \pm 0.13$ | -8.801 |
| 3 | c1ccc2c(c1)[nH]c1cccc(N3CC[NH2+]CC3)c12 | 0.839 | 0.829 | 3.187 | -2.919 | 0.679 | $-23.67 \pm 0.22$ | -11.042 |

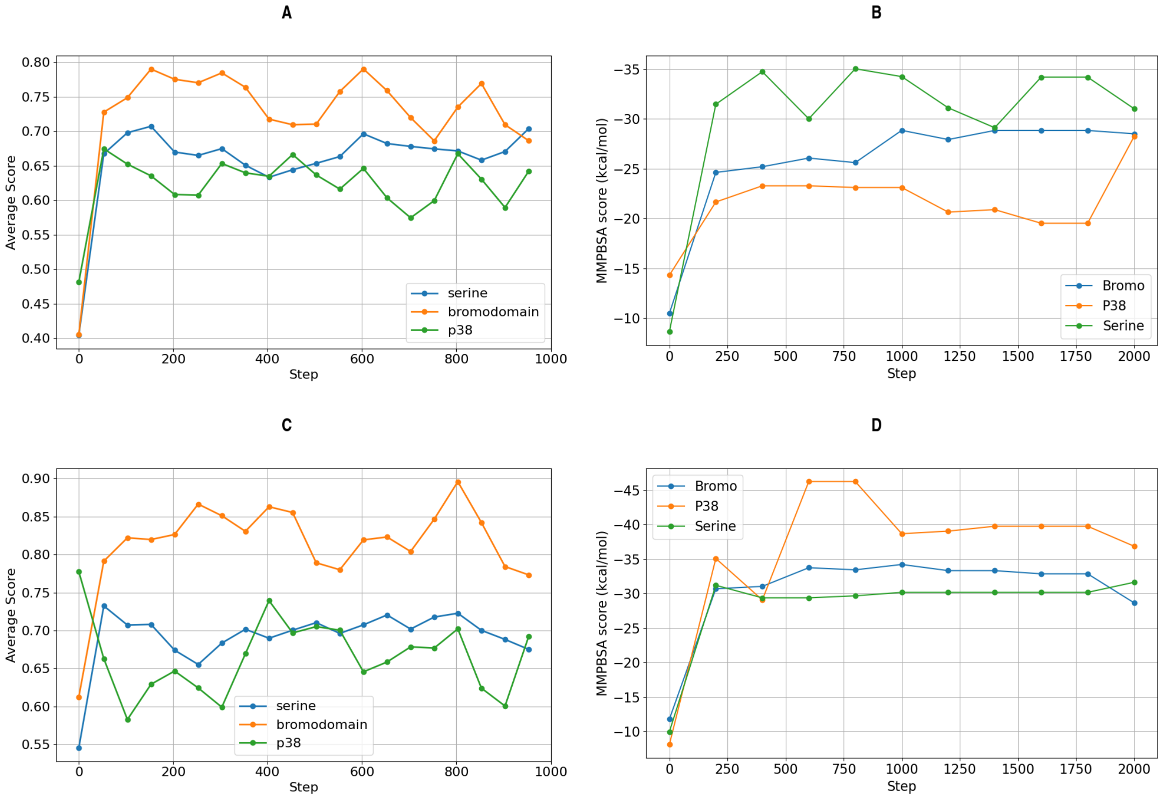


Figure SI 15: The development of the score shown for one attempt on each protein for the fragment based simulation with random and specific fragments. (A)The average of the score for every fifty steps during the fragment based simulation with random fragments for one attempt on each protein. The average was taken as running windows over 100 MC steps.(B) The development of the average MMGBSA score of the three attempts for each protein during all 2000 steps of the fragment based simulation with random fragments. (C) The average of the score for every fifty steps during the fragment based simulation with specific fragments for one attempt on each protein. The average was taken as running windows over 100 MC steps.(D) The development of the average MMPBSA score of the three attempts for each protein during all 2000 steps of the fragment based simulation with specific fragments.
